# Heterogeneous single-cell dynamics support stable population codes for objects in the mouse anterior cingulate cortex

**DOI:** 10.64898/2026.02.06.704307

**Authors:** Lucie A. L. Descamps, Wesley P. Clawson, Miguel M. Carvalho, Thomas Rogerson, Omer Hazon, Oscar M. T. Chadney, Mark J. Schnitzer, Clifford G. Kentros

## Abstract

Remembering object locations is crucial for survival, yet how the anterior cingulate cortex (ACC) encodes spatial features across repeated experiences has not been fully characterized. Using longitudinal calcium imaging in freely moving mice, we tracked excitatory ACC neurons while animals explored objects across multiple days. We demonstrate that the ACC employs a highly dynamic coding strategy: while the overall proportion of object-responsive neurons remains constant across sessions, the specific identities of these cells fluctuate, showing a continuous turnover alongside a small, stable core. This dynamic coding is modulated by behavior, with high-exploring mice exhibiting greater cellular stability. Crucially, population-level analyses reveal that stable spatial representations emerge from collective dynamics rather than fixed single-cell identities. Population decoding demonstrates that information becomes linearly separable and highly efficient at a coarser ensemble scale. Thus, the ACC achieves representational stability through emergent network organization despite persistent single-cell dynamics.

## Introduction

The anterior cingulate cortex (ACC) has been implicated in diverse cognitive processes, including attention ^1^, time perception ^2–6^, decision-making ^7^, and remote memory consolidation ^8–12^. While much of the evidence linking ACC to memory stems from studies using aversive paradigms, such as contextual fear conditioning, the strong emotional valence of such stimuli may engage specialized neural circuits ^13^, potentially limiting generalizability to neutral or positive memory representations. This raises the question of whether the ACC supports memory for non-aversive experiences, such as object exploration, and how such representations are maintained over time. Objects provide a clear avenue for studying neutral episodic-like memory in rodents ^14^, as they can be associated with spatial and temporal context without confounding emotional valence such as fear. Prior work has shown that ACC neurons exhibit firing fields near both current and previous object locations ^15,16^, suggesting a role in tracking object-related information across time. However, how these object-correlated activity patterns are specifically maintained remains to be investigated. Previous work has identified that when animals perform operational tasks, the ACC exhibits high single-neuron instability whilst maintaining ensemble-level representations ^2,17^. Whether this is a general feature of the ACC during the preservation of neutral episodic memory, or if object-related codes are stably maintained over time has yet to be tested. In other brain regions, both stable coding ^18^ and dynamic reconfiguration are found. Indeed, repeated exposure to identical stimuli or environments often yields shifting neural representations known as representational drift. This has been observed not only in multimodal association areas like the hippocampus ^19–21^ but also in primary sensory cortices, including visual ^22^, auditory ^23^, somatosensory ^24^, and olfactory ^25^ regions. As these areas provide major afferent inputs to the ACC ^26^, and given the previous evidence on ACC dynamics during action representation, it is plausible that ACC object representations also undergo dynamic remapping over time. This potentially challenges traditional engram theories that posit specific cells as the physical substrate for a memory, if the ACC supports long-term memory but its networks undergo continuous modifications rather than being a static population code. Therefore, with the methodological means to longitudinally record from the same subset of neurons, we performed calcium-imaging in freely moving mice repeatedly exposed to the same two objects in a circular arena. By tracking the same neuronal ensembles across days and analyzing both single cell and ensemble representations, we examined how object-related activity patterns emerge and evolve in the ACC. Our approach enables direct assessment of representational stability in a cortical region implicated in higher-order cognition, offering insight into how neutral episodic-like memories are encoded and distributed in the brain.

## Results

### Tracking the same cells in anterior cingulate cortex over repeated objects exploration

Using miniaturized head-mounted fluorescence microscopes (“miniscopes”), we performed calcium-imaging in the anterior cingulate cortex of transgenic animals expressing the calcium indicator GCaMP6s under the CamKII promoter (Ck2/tTa +/-; tetO-GCaMP6s +/-) (Figures 1A, S1A). Animals first explored an empty environment (Open Field sessions, Day 0). On Day 1, we introduced into the environment two objects of similar size, but with different shapes and materials, whose positions within the environment remained fixed throughout the experiment. Previous work has characterized the wide variety of ACC neurons response during the displacement of objects, with some cells tracking either the current object locations, the former locations, or a combination of both ^15^. To avoid these complexities and focus purely on the dynamics of a single experimental condition, we instead kept the identity of the objects and their locations constant throughout the experiment to specifically investigate how the same subset of ACC neurons respond to an unchanged environment. We followed the original experimental timeline from Weible et al. ^16^, such as mice had 2 object exploration sessions on Day 1, each lasting 30 minutes and separated by a 5-minute break outside the environment. From Day 2 to Day 7, animals had one 30-minute session per day, totaling 8 object exploration sessions (Figure 1B). Due to poor exploration, we discarded the data from the second object exploration on Day 1, and only kept one session per day (Figure S1G), referred to as Day 1 - Day 7.

**Figure 1.**
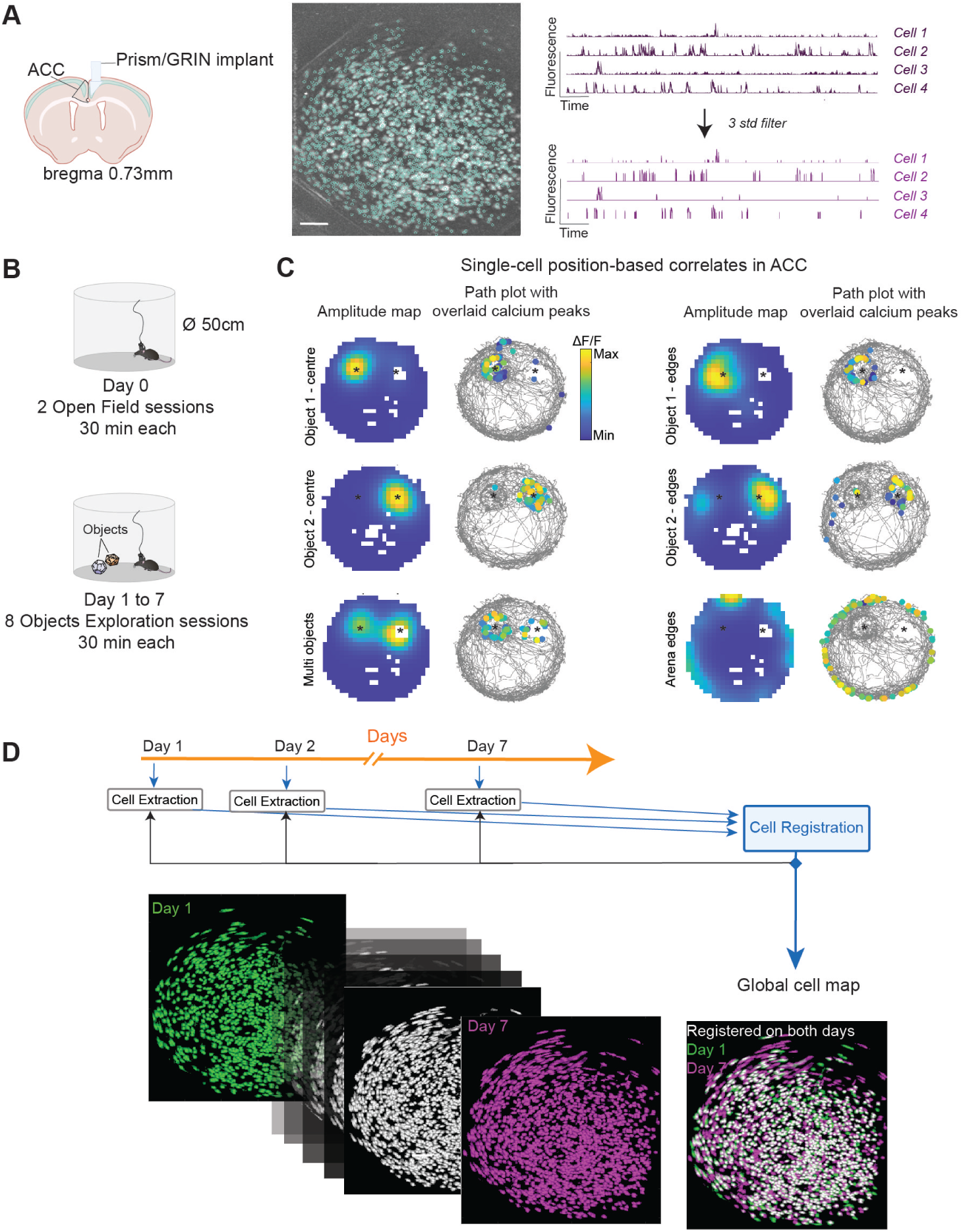
Tracking the same cells in anterior cingulate cortex (ACC) over repeated object exploration. **A)** Left: ACC was optically accessed using a Prism/GRIN assembly in mice expressing a genetically encoded calcium indicator. Middle: Representative field of view. Right: Putative cells and their fluorescence time-series were extracted from the miniscope FOV and filtered. **B)** Schematics of the behavioral apparatus during open-field sessions (top) and objects exploration sessions (bottom).**C)** Single-cells position based correlates are computed in the form of activity maps, revealing cells with preferred activity fields around objects and the edges of the arena. Stars denote the object’s locations. Examples are from mouse 7 on Day 2 (Object 1 centre, Object 1 edges, Multi objects, Arena edges) and on Day 3 (Object 2 centre, Object 2 edges). **D)** Overview of TRACKER, an iterative cell extraction and registration algorithm, used to align cells across days and track their activity during the different sessions of the experiment.

To match cells across the different days of the experiment, we used TRACKER, an algorithm performing iterative cell extraction and registration ^27^ (Figures 1D, S1B-F). Briefly, TRACKER extracts the spatial and temporal footprints of putative cells before registering them, e.g. aligning them throughout the sessions, before re-feeding the output to the cell extraction step. After iterative steps, the outputs are the spatial footprints, the fluorescence time series, and a global cell map that identifies the cells throughout the sessions (see **Methods**). We combined the fluorescence time series with the position data to create activity maps (Figures 1C, S1I) and identified localized fields either at one or both objects, around or on top of objects, and along the edges of the arena which is consistent with what has been reported previously ^16^. In our subsequent analysis, we only considered cells positively modulated by the objects if they exhibited a statistically significant spatial field at one or both object locations compared to the rest of the environment. We will refer to them as object-location cells from here on (see **Methods** for the full qualification criteria of a cell as an object-location cell). Combined with longitudinal tracking of the same cells, this enabled us to probe the dynamics of object-location coding in ACC.

### Dynamic single-cell coding underlies stable object representation across days

We first investigated whether the proportion of object-location cells in the recorded population varied throughout the experiment. As one might expect some inherent firing in any part of the environment independently of the objects ^28,29^, we used the two Open Field sessions (OF1 and OF2) to establish a baseline proportion of firing in the spatial bins that would later be the objects’ location. We observed a large increase in the proportion of object-location cells when the objects were introduced in the environment on Day 1, as can also be seen in the maps of the population stack (Figure 2A, Figure S2A). This proportion stayed stable throughout the remainder of the experiment, indicating that a similar number of cells was allocated an object coding function, rather than the recruitment of more cells (for statistical comparisons between sessions, see Table S1). Throughout the experiment, the majority of object-location cells were tuned to a single object, as opposed to both objects (Figure S3A). To further differentiate between object-location coding and pure spatial coding, we additionally analyzed two spurious object locations (Figure S2B), mirrored from the real object zones: we observed no changes in the proportion of spurious object-location cells throughout the experiment (Figure S2C). Together, these results indicate that whilst the cingulate cortex maintains a proportion of cells firing in the spurious object zones, the introduction of objects selectively recruits a larger, stable population of cells dedicated to object-location encoding.

**Figure 2.**
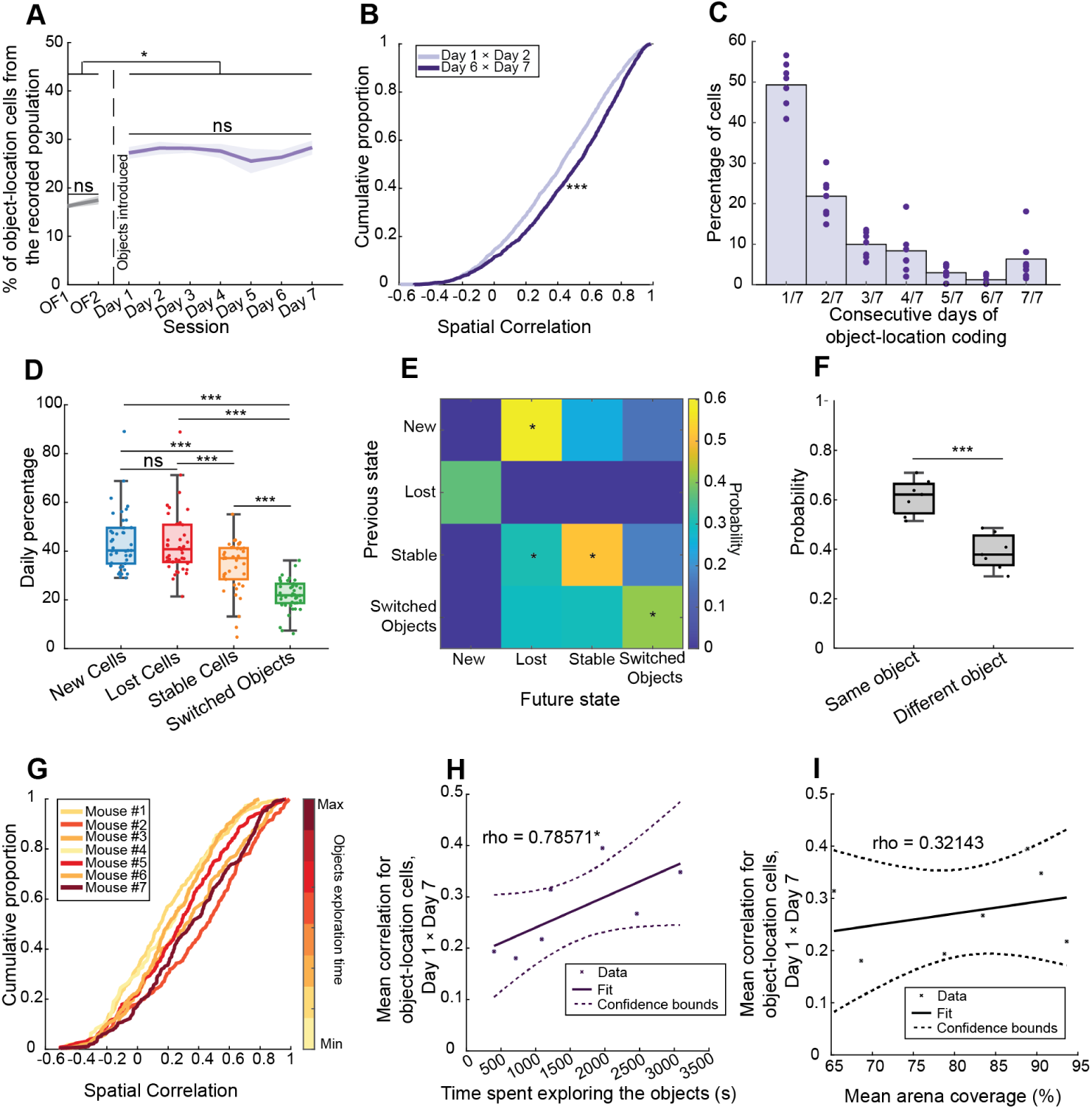
Dynamic single-cell coding underlies stable object representation across days. **A)** Proportion of object-location cells across the Open Field sessions (OF1 and OF2, grey, control sessions) and throughout repeated familiarization to the objects (Day 1 to Day 7, purple), N = 7 mice. Data are represented as mean ± SEM (dark line = mean, shaded area = SEM). For statistical comparisons between sessions, see Table S1. **B)** Spatial correlation score for object-location cells at the start (Day 1 × Day 2, light purple) and the end of the familiarization to the objects (Day 6 × Day 7, dark purple). N = 7 mice. Two-sample Kolmogorov-Smirnov test, *\*\*\*p* = 6.1090e-10. **C)** From the subset of unique object-location cells registered throughout all object exploration sessions (n = 1861 across 7 animals), percentage of object-location cells coding consecutively for the same object, out of the 7 days of the experiment. Only 6.32% of cells coded for the same object throughout the 7 days of object exploration. One dot = percentage for one animal. **D)** Percentages of new, lost, stable cells and cells switching their object preference in the object-location cell ensemble between day X and day X+1. One dot = percentage between 2 consecutive sessions from Day 1 to Day 7 for a single animal, N = 7 mice. Pooled data are represented as median and interquartile range ± nonoutlier maximum/minimum. For statistical comparisons between groups, see Table S2. **E)** Probabilities of transitioning between categories between day X and day X+1, calculated post-hoc from 2D. The star denotes a probability above chance level. **F)** For cells specifically leaving the object-location cell ensemble then joining it in a later session, probability to code for the same or a different object as they previously did. One dot = probability for a single animal. Pooled data are represented as median and interquartile range ± maximum/minimum. Two-sample t-test, *\*\*\*p* = 8.4752e-05. **G)** Spatial correlation score for object-location cells in Day 1 between their Day 1 and Day 7 fields for each animal, color-coded by the time spent exploring the two objects throughout the experiment. **H)** Correlation between the mean spatial correlation score for object-location cells in Day 1 between their Day 1 and Day 7 fields, and the time the animal spent exploring the two objects throughout the experiment. Spearman’s rho correlation, rho = 0.78571, *\*p* = 0.0167. **I)** Correlation between the mean spatial correlation score for object-location cells in Day 1 between their Day 1 and Day 7 fields, and the mean arena coverage of the animal. Spearman’s rho correlation, rho = 0.32143, *p* = 0.4976. *\*p* <0.05, *\*\*p* < 0.01, *\*\*\*p*<0.001

We hypothesized that if an animal was becoming more familiar with the environment and objects, the object-location code would be more stable and the activity fields show increased similarity throughout the experiment. To verify this, we computed the spatial correlation score between the last two days (Day 6 x Day 7) and the first two days (Day 1 x Day 2) and found that indeed the spatial correlation was higher between Day 6 and 7 than between Day 1 and 2 (Figure 2B, Two-sample Kolmogorov-Smirnov test, *\*\*\*p* = 6.1090e-10; for all pairs of consecutive days, see Figure S2E). This could suggest an increased stability of the population of object coding cells following repeated experience of the objects and the environment, though it might be a reflection of the time spent exploring the objects as it was more similar between Day 6 and 7 than Day 1 and 2 (Figure S1H, Two-sample t-test with Bonferroni correction; Day 1 vs Day 2 *\*\*p* = 0.0044; Day 6 vs Day 7 *p* = 0.9104). We observed a similar increase in spatial correlation score for the spurious object-location cells (Figure S2D, Two-sample Kolmogorov-Smirnov test, *\*\*\*p* = 3.9947e-13; for all pairs of consecutive days, see Figure S2F) meaning that irrespective of the presence of an object, spatial tuning stabilizes over experience, consistent with previous reports of regularization over time ^25,30–32^.

Given that the object-location cells’ field stability increased with exposure, we further asked if the acuity of an object-modulated field is predictive of its stability across sessions, as this would imply that the neurons are “tightening” their tuning curves to stabilize a given field. Using a template-matching procedure ^33^, we investigated the relationship between each object-location cell’s object-tuning score and the stability of its field. We found that the object-tuning score on day X held a very low predictability of the stability of the cell’s activity field between day X and X+1, as there was only a weak correlation between the object score on day X and the spatial correlation score between day X and X+1 (Figure S3B, Spearman’s rho correlation, rho = 0.15, *\*\*\*p* = 1.4592e-87). This showed that even cells with very sharp activity fields around the objects could display a different activity pattern upon the reintroduction to the environment and the objects. Overall, we found that the vast majority of cells that were object-location cells on Day 1 were not stable, with nearly half of them not coding for the same object in the next session, and progressively fewer coding for increasingly longer number of consecutive days. The exception was a small population of object-location cells coding for the same object location in every session (Figure 2C, 6.3% of cells out of 1861 unique object-location cells on Day 1 registered in all subsequent sessions), suggesting the possibility of a stable set of “core cells”. This over-representation of object location cells that were stable throughout the experiment was not found using spurious object zones (Figure S2G, 1.34% cells out of 823 unique spurious object-location cells on Day 1 registered in all subsequent sessions).

Despite a stable proportion of object-location cells across days (Figure 2A), the object-tuning score poorly predicted the field stability, and a small number of cells reliably coded for the same object in every day of the experiment (Figure 2C).

Next, we hypothesized that the object-location cell ensemble (that is, the population of object-location coding cells) would drastically change from one session to the next. To that end, we restricted our next set of analysis to cells that were registered throughout all sessions with objects in the environment (Day 1 to Day 7, average number of cells 571±195, N = 7 animals). We questioned whether these cells, active everyday, would have the same behavior - code for an object or not.

We postulated that on any given day (day X), the object-location cell ensemble - corresponding to all cells with a significant spatial field at one or two objects - would comprise cells (1) that were already object-location cells the day before (day X-1) for the same object (“Stable cells”), (2) cells that were already object-location cells the day before but for a different object (“Switched object”) and (3) cells that were not object-location cells the day before (“New cells”). Additionally, there will be (4) cells part of the object-location cells ensemble on day X, but not day X+1 (“Lost cells”). We computed the percentages of cells falling into these four categories on a daily basis between day X and day X-1 for Stable, New, and Switched object-location cells; and between day X and day X+1 for Lost cells. We found that there was an equivalent, nearly matched proportion of New (median = 40.27%) and Lost (median = 40.82%) cells, meaning there was as much inflow as outflow of cells in the object-location cell ensemble. A smaller but non-negligible proportion of cells exhibited a stable object code between two consecutive days (median = 37.14%), whilst fewer cells retained an object-location coding function but for the other object-location (median = 21.92%) (Figure 2D, for statistical comparisons between groups, see Table S2). Importantly, for every object-location cell falling into these categories, we performed a post-hoc quantification of the matched centroids’ location distances to check if the inflow/outflow dynamics we uncovered could be an artefact of the cell registration algorithm. We found that, on the contrary, the distances were very small (mean distance “New” cells 2.59µm; mean distance “Lost” cells 2.76µm; mean distance “Stable cells” 2.52µm; mean distance “Switched object” cells 2.47µm; Figure S3C-F). As such, the dynamics described were not because of centroids wrongly considered as the same cell across days.

To describe the relationship between single cells and ensemble dynamics further, we computed the post-hoc probabilities for cells to transition from one category (“Previous state”) to another (“Future state”) (Figure 2E). Some transitions were impossible, for example a Lost cell could only be New. We compared this to uniform probabilities (see **Methods** and Figure S3G) and found that the following happened above chance level: “New” cells were more likely to exit the object-location cells ensemble, “Stable” cells were most likely to keep coding for the same object (though the probability to exit the ensemble was also slightly higher than chance), and “Switched” cells tended to switch their object preference again in the next session.

We also further investigated the cells transitioning from Lost to New - meaning object-location cells that left the object-location cell ensemble, then entered it again - and found that significantly more cells coded for the same object as previously, rather than the other object (Figure 2F, Two-sample t-test, *\*\*\*p*= 8.4752e-05); or for the first object they ever coded for (Figure S3H, Two-sample t-test, *\*\*\*p* = 5.0151e-10).

We next wondered if the dynamics described for the object-location cell ensemble would be similar for purely spatial coding. To that end, we performed a similar analysis with the spurious object zones (Figure S2H). Interestingly, we found again a nearly-matched proportion of cells in “New” and “Lost” categories (median spurious location “New” = 67.39%, median spurious location “Lost” = 64.45%), implying that a steady inflow/outflow is a strategy employed by the cingulate cortex to maintain a stable population representation of both spurious and object-location firing. However, there was a smaller proportion of “Stable” spurious object-location cells than real object-location cells (median spurious location “Stable” = 15.14%, median real location “Stable” = 37.14%), suggesting that the presence of an object increases the likeliness of a cell to remain tuned to that location. So far, our results demonstrate a fine balance between stable and dynamic coding at the single-cell level, despite the animals exploring an unchanged environment.

Since the mice exhibited different degrees of interest and exploration of the objects (Figure S4A), we sought to explore the link between exploratory behavior and the stability of the object-location neural code. To do this, we quantified the spatial stability of object-location cells between Day 1 and Day 7. For each animal, we plotted it as a cumulative proportion and color-coded it by the degree of exploratory behavior throughout the experiment, using the total time spent exploring the objects (Figure 2G) and observed a clustering by exploratory behavior. We further quantified the relationship between behavior and the stability of the object code and found that the stability of the neural code was positively influenced by the time spent exploring the objects (Figure 2H, Spearman’s rho correlation, rho = 0.78571, *\*p* = 0.0167), but not by the coverage in the arena (Figure 2I, Spearman’s rho correlation, rho = 0.32143, *p* = 0.4976). This suggests that the spatial stability was not modulated by overall exploratory behavior in the environment.

We wondered whether this result was simply a reflection of the time the animals spent sampling the objects on Day 1 and Day 7. To answer this, we plotted the mean spatial correlation score of object-location cells between Day 1 and Day 7 against either the time spent exploring the objects on Day 1 (Figure S4B), the time spent exploring the objects on Day 7 (Figure S4C), or the time spent exploring the objects between Days 1 and 7 (eg. from Day 2 to Day 6, included). We found a slight, non-significant anti-correlation between the stability of the neural code and the exploration of Day 1 (Spearman’s rho correlation, rho = −0.17857, *p* = 0.7130) and a small, non-significant correlation with the exploration on Day 7 (Spearman’s rho correlation, rho = 0.2857, *p* = 0.5559. However, there was a strong, significant correlation with the time spent exploring the objects between Day 2 and Day 6 (Spearman’s rho correlation, rho = 0.78571, *\*p* = 0.0480), indicating that a bias in the objects exploration on either Day 1 or Day 7 is not driving the stability of the spatial code, but it is rather the exploration (and potentially, interest) throughout the experiment. Taken together, these results suggest that the stability of the code was not merely influenced by how much an animal was exploring its environment, but by the amount of exploratory behavior around the objects in the environment. This implies that ACC neurons do not represent a mere spatial map of the environment, but instead could integrate task demands and internal state.

### Cell assemblies constructed with coarse-graining highlight environment features

We have now characterized the dynamics of single-cell object-location coding. Given the extensive single-cell turnover and despite the intrinsic biological “noise” observed at the single-cell scale ^34^, we asked whether stability emerges at a higher level of organization. To address this, we moved to a coarser-grained representation that captures collective dynamics across neural ensembles, allowing us to investigate population-level stability and structure beyond the variability of individual units.

We chose to use recursive coarse-graining ^35^ to construct cell assemblies (Figures 3A-B, S5A-B) because it enabled us to perform population analyses on multiple scales. Briefly, we first identified the two most correlated neurons in the FOV, summing and normalizing their activity trace, to create a new signal (“coarse-grained signal”, CS). We then removed these two neurons from our pool of cells and looked for the next two most correlated neurons, and repeated the procedure until all neurons had been paired: this was defined as a coarse-graining step (or clustering step). We iteratively performed a new coarse-graining step until no more signals could be paired together (for more details, see **Methods**).

**Figure 3.**
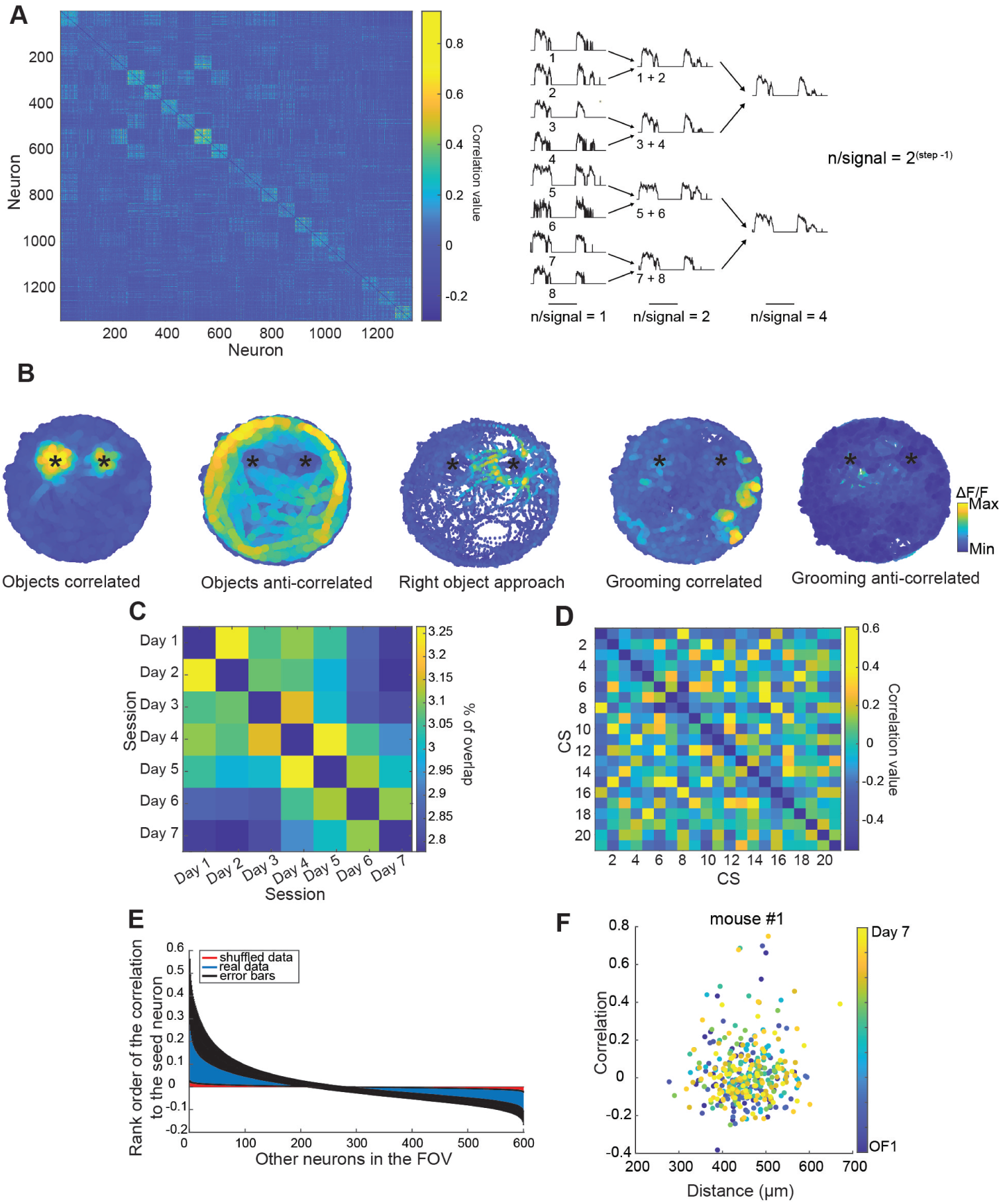
Cell assemblies identified with coarse-graining highlight environment features. **A)** Left: Correlation matrix of single cells recorded in an example session. For visualization purposes, the values on the diagonal where a neuron is correlated to itself have been replaced by NaNs. Right: Schematics illustrating the coarse-graining procedure. **B)** Examples of coarse-grained signals (CS) obtained from step 7 (64 neurons per signal), displayed on the animal’s path plot. Assemblies can be anti-correlated with each other, such as the two on the left. Some assemblies display more than just object-location information, such as the one in the middle whose activity is ramping up as the animal is approaching the right object (object 2); or the two on the right that are modulated by grooming behavior. The stars denote the objects’ locations. **C)** Matrix visualizing the percentage of unique cells overlapping in CS between days. For visualization purposes, the values on the diagonal where the overlap would be 100% have been replaced by NaNs. **D)** Correlation matrix of CS obtained from step 7 in an example session. Some signals are highly correlated with each other, which means they will be coarse-grained together in the next step. Similar to A), the values on the diagonal where a CS is correlated to itself have been replaced by NaNs. **E)** Rank order correlation between a random seed neuron and other neurons in the FOV, highlighting correlation and anti-correlations between neurons. Error bars represent standard deviation. **F)** Scatter plot of the correlation values between 2 neurons and their anatomical distance in the FOV, at step 7, for all experimental sessions for a single mouse. For each session and each mouse, we computed the slope of the correlation between these 2 metrics and plotted it in Figure S6B.

At each iteration, or step, the number of neurons composing each coarse-grained signal was 2^(step-1)^ (Figure 3A): this means that whilst the number of neurons per coarse-grained signal was fixed at every step, the number of coarse-grained signals produced at each iteration was dependent of the number of cells recorded in that session. Since we repeated the coarse-graining procedure on every session, and because the neurons were grouped together based on their temporal activity pattern in that given session, this also means that the assemblies we constructed were composed of different cells from one session to the next, even at a similar step. For this reason, and in line with our results from Figure 2 demonstrating that single-cells are mostly exhibiting dynamic activity patterns across days, we found only a very small proportion of unique cells being coarse-grained together on different days (less than 5%, Figure 3C). After obtaining our coarse-grained signals at different steps, we could plot their activity over the path plot of the animal (similarly to what we have done previously with single-cell activity traces) to examine assembly maps ^36^. For visualization purposes, we will use step 7 (K7), which corresponds to 64 neurons per coarse-grained ensemble (Figure 3B).

We found many position-based signals highlighting different features of the environment such as the edges of the arena and the objects (see Figures S6G-H for proportions of object-location coding assemblies), including more complex patterns such as activity ramping up to an object or correlated with non-spatial behaviors such as grooming (see Figure S5C for additional characterization of grooming-correlated activity). Some signals were strongly anti-correlated with each other. These anti-correlation patterns at the population level between coarse-grained signals could be recapitulated at the single cell level, when selecting a random seed neuron and ranking the correlations between that neuron and all the other neurons in the FOV (Figure 3E). As coarse-graining clusters cells based on their degree of correlated temporal activity, we checked whether neurons anatomically close to each other in the FOV would be clustered together. We found no correlation between the anatomical distance between neurons and their levels of co-activity, neither at different clustering steps, recording days, or animals (Figures 3F, S6A-E). We then hypothesized that the repeated exposure to the environment and the objects could lead to a change of the degree of correlation between neurons. To that end, we extracted the top 10% correlation values between single cells after one step of coarse-graining and compared them between Day 1 and Day 7. We chose to look at only the top 10% values, instead of all the correlation values, as we wanted to avoid any potential difference being drowned out by the mean. We found no difference between Day 1 and Day 7: single-cells whose activity correlated together did so at the same degree at the start and end of the experiment (Figure S6F).

We next sought to investigate how core cells (object-location cells coding in every session for the same object, see Figure 2C) related to coarse-grained signals. Given that CS were for the majority not composed by the same cells between days (Figure 3C), would core cells on the contrary cluster in the same assemblies? We found that on average, 50% of core cells would overlap and be coarse-grained in the same CS (Figure S6I). Surprisingly, even core cells from different objects were clustered together (Figure S6J: CS #7 and CS #14 both comprised core cells from object 1 and object 2). This suggests that the activity of some core cells might be modulated by more than the object identity and location.

### Decoding performance evolves with experience and is optimal using population data

Coarse-graining correlated neural activity revealed ensemble-level structure, where the joint activity of neurons formed clear and stable representations of environmental features and ongoing behavior, despite substantial turnover at the single-cell level.

We sought to verify this and explored how this evolved with experience using a decoder of neural activity to identify where in the environment the mouse was situated (Figure 4A). We first constructed regions of interest (ROIs) by applying a Gaussian blur, and by overlaying the position path on it we created position labels of the animal in the environment (Figure S7A, see **Methods**). We used decision tree classifiers to decode the animal’s position (see **Methods**). This approach provided two interpretable metrics: (1) the classifier’s decision output, which allowed us to compute decoding accuracy, and (2) the tree’s maximum depth, which reflected the maximum number of decision steps required to reach a classification (Figure 4B). For each animal, we ran the decoder on all possible coarse graining steps, trained on both real and shuffled data (see **Methods**).

**Figure 4.**
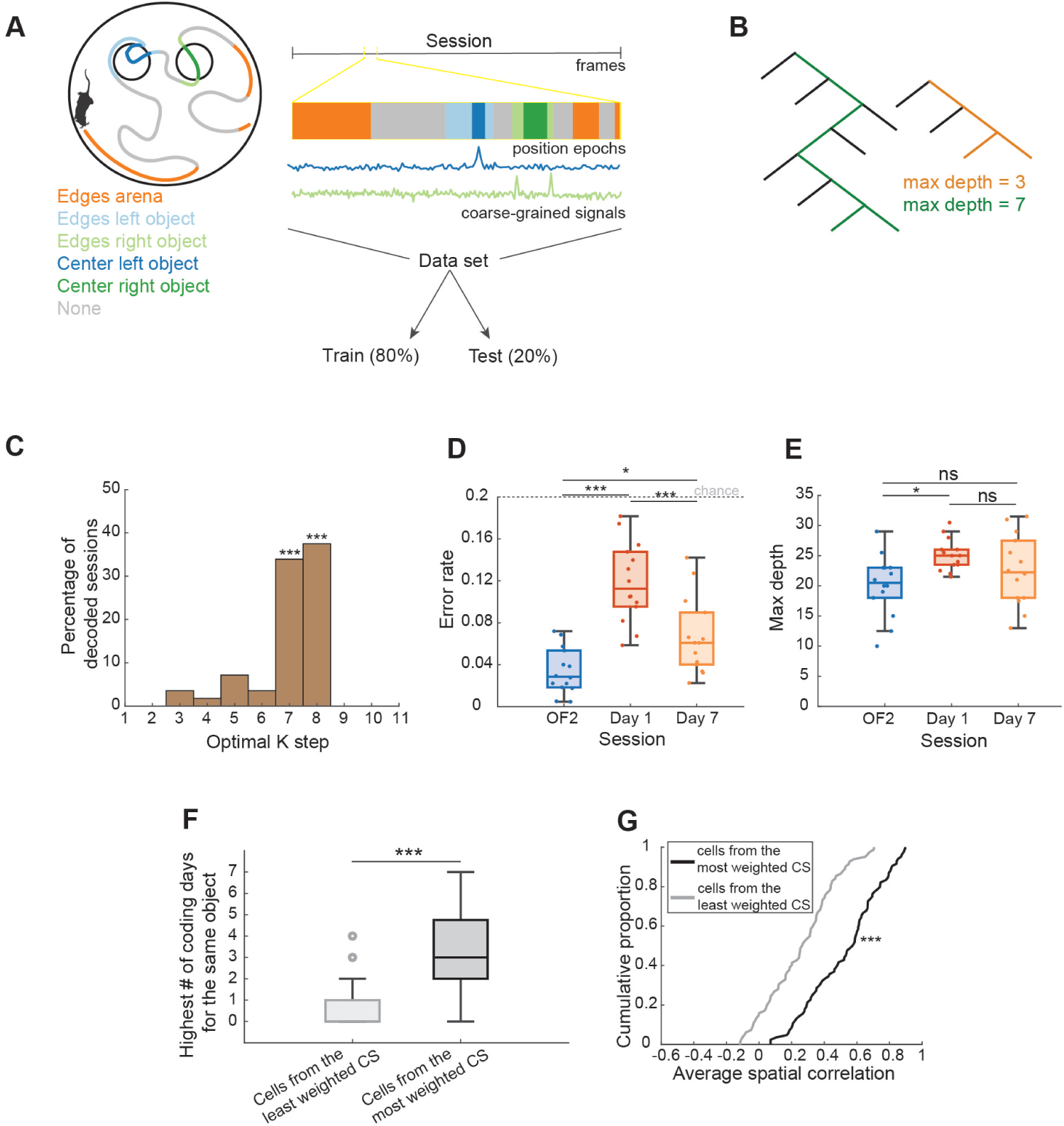
Decoding performance using decision trees evolves with experience and is optimal using population data. **A)** Schematics illustrating the decoding procedure. Position labels are created from 5 different zones explored by the animals. Coarse-grained signals are used to decode the position of the mouse within the environment. Training and decoding are performed within a single session. **B)** Schematics illustrating two decision trees of different depths. The tree on the left (green) has a maximum depth of 7, whilst the tree on the right (orange) has a maximum depth of 3, meaning that the orange tree needs less decisions than the green one before outputting a result. **C)** Optimal ensemble size for decoding position data. N = 7 animals. The decision trees were most efficient and accurate when using population data from K7 (64 neurons per ensemble, Z-test, Z-statistic = 4.15, *\*\*\*p* = 3.2858e-05) and K8 (128 neurons per ensemble, Z-test, Z-statistic = 4.9193, *\*\*\*p* = 8.6832e-07). **D)** Error rate at K7 and K8 when decoding position labels in the last open field session (OF2), the first session with objects (Day 1) and the last session with objects (Day 7). One dot = value of error rate for a single animal at either K7 or K8, as such there are 2 dots per animal for a given session. N = 7 mice. ANOVA, OF2/Day1 *\*\*\*p* = 1.061e-07; OF2/Day7 *\*p* = 3.09e-02; Day1/Day7 *\*\*\*p* = 4.37e-04. **E)** Maximum tree depth at K8 when decoding position labels in the last open field session (OF2), the first session with objects (Day 1) and the last session with objects (Day 7). Similar to D), one dot = value of maximum depth for a single animal at either K7 or K8, as such there are 2 dots per animal for a given session. N = 7 mice. ANOVA, OF2/Day1 *\*p* = 1.87e-02; OF2/Day7 *p* =3.59e-01; Day1/Day7 *p* = 3.19e-01. **F)** Highest number of coding days for the same object throughout the experiment for cells picked from the least weighted CS (light gray) and most weighted CS (dark gray) when decoding position labels on Day 1 at K8. N = 91 cells from the least weighted CS pooled from 7 mice; N = 91 cells from the most weighted CS pooled from 7 mice, Two-sample t-test, *\*\*\*p* = 1.2455e-27. **G)** For cells picked from the least weighted CS (light gray) and most weighted CS (dark gray) when decoding position labels on Day 1 at K8, average spatial correlation across the experiment. N = 91 cells from the least weighted CS pooled from 7 mice; N = 91 cells from the most weighted CS pooled from 7 mice, Two-sample Kolmogorov-Smirnov test, *\*\*\*p* = 7.9649e-09. In D-F, pooled data are represented as median and interquartile range ± nonoutlier maximum/minimum. *\*p* <0.05, *\*\*p* < 0.01, *\*\*\*p*<0.001

Since coarse-graining allowed us to query the population at multiple scales, we first used the decoder results to investigate which coarse-graining step, or assembly size, was the most efficient for position-based coding in ACC. We combined both outputs from the decision trees and assumed that the optimal performance would be a combination of minimum error rate and minimum depth. For each mouse and each session, we normalised the error rate and maximum depth values, summed, and checked at which K step we would find the lowest values. We found that K steps 7 and 8, respectively ensemble sizes of 64 and 128 neurons, appeared to be the optimal clustering steps for decoding position-based labels (Figure 4C, K7: z-test, z-statistic = 4.1527, *\*\*\*p* = 3.2858e-05; K8: z-test, z-statistic = 4.9193, *\*\*\*p* = 8.6832e-07, Figures S7B-C), demonstrating effective compression of the single-cell code.

We next hypothesised that the performance of the decoder would vary when the environment explored by the animals was subjected to changes or becoming over familiar. To answer this, we compared the error rate and maximum depth from K7 and K8 at the following sessions: Open Field 2 (OF2), Day 1 and Day 7 (Figures 4D-E, see Figures S7D-E for all sessions as well as Tables S10 and S11 for statistical comparisons).

Between OF2 and Day 1, which corresponded to the introduction of the objects in the environment, the error rate significantly increased (median OF2 = 0.0292, median Day 1 = 0.1125, ANOVA, *\*\*\*p* = 1.061e-07) as did the maximum depth (median OF2 = 20.5, median Day 1 = 25, ANOVA, *\*p* = 1.87e-02), which means the overall decoder performance worsened when the content of the environment changed.

Between Day 1 and Day 7, the error rate decreased significantly (median Day 1 = 0.1125, median Day 7 = 0.0609, ANOVA, *\*\*\*p* = 4.37e-04), such that the difference in error rate between OF2 and Day 7 was minimal (median OF2 = 0.0292, median Day 7 = 0.0609, ANOVA, *\*p* = 3.09e-02). The maximum depth was not significantly different between Day 1 and Day 7 and OF2 and Day 7 (median Day 1 = 25, median Day 7 = 22.25, ANOVA, *p* = 3.19e-01 between Day 1 and Day 7, *p* = 3.59e-01 between OF2 and Day 7). Taken together, these results show that whilst an abrupt change in the content of the environment (i.e. the introduction of the objects) reorganised the population representation, thus worsening the decoding performance, it was dynamic enough to lower the error rate again given more exposure to the environment.

We then checked whether stable single-cells (see Figure 2) were more likely to support the decoder performance at the optimal ensemble size. For each animal, we extracted the coarse-grained signals that had the most and least weights in the classification tree on Day 1 for K8 (see **Methods**). We then isolated for both most weighted and least weighted signals the single-cells whose activity were most correlated with the signals themselves, assuming these single-cells may play a critical role in their respective ensembles. We specifically chose to look at Day 1 - the first day with objects - to assess how these cells would code in the remaining days of the experiment. For each of these single-cells, we computed the highest number of days they would code for the same object - if any - and their average spatial correlation score across days. We found that cells from the most weighted ensemble on Day 1 would develop across the experiment a significantly more stable object code and a higher average spatial correlation score, than cells from the least weighted ensemble (Figure 4F, median cells from most weighted CS = 3, median cells from least weighted CS = 0, Two-sample t-test, *\*\*\*p* = 1.2455e-27; Figure 4G, median cells from most weighted CS = 0.5772, median cells from least weighted CS = 0.2660, Two-sample Kolmogorov-Smirnov test, *\*\*\*p* = 7.9649e-09).

We showed previously that using coarse-graining, we could construct cell assemblies and reveal correlates not only for objects but also behaviors such as grooming (Figures 3B and S5C). We therefore asked what is the relationship between position-based and behavior codes in the cingulate cortex. If the optimal ensemble sizes for decoding position-based labels are 64 and 128 neurons, does that mean it will generalize to any population code in the ACC, irrelevant of the task performed, or are these ensemble sizes specific to correlates that are position dependent? We sought to answer this question by running the decoder on behavior-based labels that would be position independent. We chose the three following behaviors: grooming, rearing and resting (see **Methods**). We followed a similar procedure as for the position-based labels, normalising the error rate and maximum depth values then summing them up to find at which clustering step we would find the lowest values, meaning a combination of minimum error rate and minimum depth. We found that again K steps 7 and 8 (ensemble sizes of 64 and 128 neurons respectively) were the most prevalent and appeared to be the optimal steps for decoding behavior-based labels as well (Figures S7F-H, K7: z-test, z-statistic = 4.1576, *\*\*\*p* = 3.216e-05; K8: z-test, z-statistic = 2.7969, *\*\*p* = 5.159e-03).

## Discussion

We used calcium-imaging to longitudinally record the activity of neurons in the anterior cingulate cortex while animals were self-engaging in the exploration of objects, over several days. Our data show that whilst there is always a subpopulation of recorded neurons that exhibit firing fields at one or both object locations, which neurons exhibit these fields remains highly dynamic. Given the role of the ACC in long-term memory and memory consolidation, this directly challenges classical engram theories which posits that specific, stable ensembles of dedicated cells are the physical substrate of a memory. Interestingly, we observed a modulation by behavior as the animals that spent the most time at the objects locations displayed a more stable object code. In addition, we examined population activity using coarse-graining and found that stable representations of environmental features and complex behaviors emerge at the ensemble level despite the dynamic nature of single-cell coding, suggesting a population-level alternative to the static cellular engram. We used decision trees to decode neural activity and showed that the representation of the animal’s position and behavior is more linearly separable at a coarser scale. We further characterized the optimal ensemble size to be in the range of 64-128 neurons, whether coding for position-based or behavior-based correlates; demonstrating a generalized coding regime in the cingulate cortex. Finally, we showed that the decoder performance on Day 1 of object exploration was most supported by single cells that would ultimately develop a stable object code, suggesting a predictability to the stability of the object code.

We have kept the objects’ identity and location identical throughout the experiment, but what we describe as “exploration” includes a wide range of behaviors (e.g. whisking, sniffing, climbing). Therefore, the object correlates we are recording are most probably a conjunction of object identity, spatial location, and behaviors elicited by object investigation, all of which could be modulated by the animals’ evolving sense of familiarity. Some of our data imply a larger role of behavior in the neural activity we are recording: we show that the stability of the object code is positively influenced by the time the animals spent investigating the objects; and that core cells from both objects can be coarse-grained together, suggesting an underlying similarity in what drives activity in these cells. Our result showing that the object score (the “tightness” of the field at the object) only holds a small predictability of the field stability might also point towards a greater influence of behavior in cingulate coding. Indeed, if the activity of a neuron is driven by a specific exploratory behavior that is not repeated at the same object, or not repeated at all in a subsequent session, then that neuron will have a low spatial correlation score.

Our findings reveal a positive correlation between the stability of object-location cell spatial fields from Day 1 to Day 7 and the amount of time animals spent interacting with objects from Day 2 to Day 6. Despite literature showing that rodents typically spend less time with familiarized objects during novel object recognition tasks ^37^, greater object-location cell stability was observed in animals that spent more time with the objects. This discrepancy may reflect the nature of our experiment, which allowed for self-directed, unbaited exploration over multiple days as a form of incidental learning ^38^; rather than explicit familiarity or reward-driven learning. These results suggest that object-location cell stability in the ACC reflects ongoing cognitive engagement and environmental salience rather than a rigid locally stored memory trace.

Indeed, interest and attention have been described as mechanisms that can positively influence the stability of neural codes, as seen for example in the CA1 region of the hippocampus: place cells are more stable in contexts where the animal has to pay attention to its environment ^39^. In this study, the environment with the lowest degree of behavioral relevance, where place cells were the least stable, was an environment where there was no task to perform, the animal was not hungry, and no food was thrown in the environment to encourage spatial exploration - a condition highly similar to our experimental setup. In this context, the cingulate cortex’s known involvement in attention ^1^, goal-tracking ^40,41^, and outcome prediction ^42,43^ supports the idea that animals continuously monitor salient cues in their surroundings. Similarly to CA1, the engagement of the animal might help “lock” correlates that could be of high behavioral relevance. Future research could explore how object code stability changes in tasks where objects have positive valence and explicit behavioral relevance, such as reward-driven paradigms that require focused attention to object identity and location.

There have been several reports that the ACC is continuously integrating ongoing internally and externally generated sensory, cognitive, and behavioral information across different timescales, tasks, and species ^44–47^. One recent study showed that in rats repeatedly performing the same behavioral task, the ACC distinctly represented each behavioral trial, but the population activity drifted such that the most recent trials were more similar than the older ones ^6^ - suggesting that the ACC is tracking accumulated experiences and separating them over time in a structured overlapping manner. The neural correlates we are observing could therefore be a tracking of features in the environment, explaining why the population is stably representing them, both for salient locations (like the ones containing the objects) and non-salient. However, the fact that it is an overlapping but distinct population coding for the objects on each day may enable accurate discrimination between memories of similar events occurring separately across time, directly challenging a static, cellular engram. In our data, this is evidenced by a consistent proportion of cells coding for object and spurious locations, supported by a precise, near-matched proportion of cells entering and leaving the ensembles coding for these locations. Alternatively, the cingulate cortex might simply be encoding what is needed in downstream regions, reflecting both current behaviors and encountered context, and the identity of the single-cells performing the encoding is irrelevant.

Is the dynamic single-cell encoding we see in our dataset a hallmark of the cingulate cortex? Previous work carried out in rats using electrophysiology has shown that across different tasks and contexts, the ACC not only exhibits high single-neuron instability, but also stable ensembles by utilizing balanced network reconfigurations ^2,17^, a mechanism very similar to our finding of near-matched proportions of single cells entering and exiting ensembles coding for spatial locations. Our data now demonstrate that this dynamic architecture also applies to object-location coding and extends across multiple days to examine the consolidation process. This can be directly challenging classical engram theories stating the locus of a memory is stored in specific cells, whose nodes strengthen during memory consolidation. However, we also report through our population analyses no difference in the degree of single-cell temporal correlations between the start and the end of the experiment. While contextual fear studies suggest ACC engrams can be stably reactivated ^48^, our observations of both turnover and partially stable object-location cell ensembles are not necessarily incompatible. For example, De Nardo et al. ^49^ carried out an elegant study in the prelimbic cortex, showing that when neuronal ensembles are tagged either during contextual fear learning or retrieval at recent time points (1 day, 7 days or 14 days after learning) and during retrieval at a remote time point (28 days), the most similar ensembles are between recent and remote retrievals. This suggests that there is both a proportion of tagged neurons stably encoding the features of the environment used in the contextual fear assay, and another population undergoing memory-dependent reorganization. Here we see potentially a very similar mechanism during objects exploration in the ACC: some neurons reliably reactivated across sessions and others involved in flexible remapping. Our decoder results suggest some predictability in which cells would become reliably reactivated across days. Specifically, single cells part of assemblies most supporting the decoder performance on Day 1 of objects exploration displayed a high average spatial correlation score and stable object code. However, this predictability was limited to a small subset of cells and did not imply the existence of a fixed or persistent ensemble supporting object memory. A provocative take on this would be to hypothesize that the cingulate cortex does not in fact hold any ensembles to support memories. Rather, its function is to map features, internally or externally generated, so that they can be read out to downstream regions - akin to a RAM or a movie screen. The mapping of these features would follow rules specific to the task or context: in our case, a small overlap of single-cells from one day to the next, but always roughly 30% of the population coding for objects.

How dynamic single-cell tuning can support stable behaviors is also of major interest in the field of representational drift, as it has long been assumed that long-lasting neural codes and stable engrams support long-term memory. The brain is not static; elements such as neurons, synapses, and dendritic spines are continually renewed or modified, introducing inevitable biological noise and instability ^50^. Recent evidence suggests that representational drift is a natural feature of memory circuits, not a failed state or noise, but a reflection of this biological dynamism ^51^. There have been several hypotheses as to why drift is observed in neural circuits in both coding and non-coding space - some posit that neural codes are constantly updating to support ongoing, hierarchical forms of learning, while redundancy within neural populations allows for robust behavior despite changes at the single-cell level ^50,52^. Biological noise, such as spontaneous synaptic fluctuations and dendritic remodeling, may actually support flexible population coding: drift in coding and non-coding dimensions can be absorbed at the population level so overall behavioral output remains stable. Theoretical work highlights how compensatory plasticity, through homeostatic or Hebbian processes, might underlie this stability ^52,53^, and representational drift could serve as a readout of large-scale plasticity mechanisms at play throughout the system ^54^. As the ACC is a highly cognitive region ^45^, it might influence the rate at which neural activity drifts. Further experiments are needed to uncover whether drift dynamics in the ACC are similar to other regions, such as recording simultaneously several areas across the brain during the same task.

While representational drift remains an open topic, our data show that stable population-level correlates exist each day, even as the specific neurons within those populations change. Rather than imposing object-based populations, we sought a method to (1) probe population coding at multiple scales, potentially revealing emergent collective features not accessible at the single-cell level, and (2) define populations based on pairwise correlative structure, given the established importance of correlations in population coding ^55,56^. Neural data are known for rich correlation structure, often indicating higher-order interactions, as demonstrated in our dataset and mirrored in correlation/anti-correlation patterns observed in visual cortex ^57^. This rationale informed our coarse-graining approach, inspired by statistical physics, where averaging uncorrelated activity yields Gaussian distributions. If single-cell correlations were trivial, recursive coarse-graining would quickly erase correlative structure across the population. While future work could examine the persistence of non-Gaussian structure in our data and further analyses into scale invariance and criticality, it is interesting that this unbiased method produced coarse-grained populations which seem to code for objects in a stable manner across days. By refraining from imposing a minimum correlation threshold, we ensured an unbiased aggregation, although this led to less functionally coherent pairs in the final steps and a rise in decoder error. This phenomenon underscores the trade-off between ensemble size and coding fidelity - optimal ensemble sizes, determined here by decoder performance, consistently fell within the range of 64–128 neurons, for both position and behavior decoders. This is in line with similar findings in the visual cortex where ensembles of 50 neurons are as efficient as 200 neurons to unravel scale-free dynamics ^57^, and motor cortex where only 45 neurons are as efficient as 200 to explain 95% of the variance ^58^. While the size of the assembly may vary with its theoretical function ^59^, minicolumns in the cortex are hypothesized to be around the size of 80-100 neurons ^60,61^. Therefore, it may be biologically or evolutionarily optimal for assemblies, such as those defined here, to fall within a similar range, which then contributes to higher-order, hierarchically organized brain function.

Our results provide evidence that object encoding in the ACC is mostly dynamic at the single-cell level whilst allowing for modulation by behavior and attention. Populations consistently code not only position-based correlates such as the objects locations and the arena edges, but also position-independent, stereotypical behaviors such as grooming. Our decoder shows that cell assemblies comprising 64-128 neurons are optimal to decode both position-based and behavior-based epochs, suggesting that the cingulate encoding regime is invariant to the task and that population codes are preferable to individual neurons as input to downstream regions. This collective readout is an alternative to a rigid, cell-specific engram, illustrating how a memory substrate can remain functional despite a dynamic single-cell code. Future work could investigate what happens when one of the 2 objects is removed 30 days after the last habituation session to determine whether cells exhibiting firing fields at the location of the now absent object (so called absent-object cells described previously ^16^) were already part of the object-location cell ensemble during the habituation sessions.

Overall, our data provide knowledge into how the cingulate cortex supports episodic-like memory for non-aversive experiences; and open avenues for investigating if population codes are scale-invariant, and how they relate to criticality and correlate with behavior.

## Limitations of the study

In our behavioral manual annotation for the decoder, we could not confirm the animals were sleeping as we lacked electrophysiological signatures of sleep, as such we used the term “resting”. Our study was also limited to female mice. We do not expect that the physiology of the anterior cingulate cortex should vary by sex, however future studies could investigate whether our findings generalize to male mice.

## ACKNOWLEDGEMENTS

We thank Tuce Tombaz for help with training in surgeries, Richard Gardner and Rafael Pedrosa for help with the synchronization between behaviour and imaging, Nick Robinson for helpful discussions and Matteo Guardamagna for comments on the manuscript. Supported by the Centre of Excellence scheme of the Research Council of Norway - Centre for Neural Computation (Grant 223262/F50) and Center for Algorithms in the Cortex (Grant 332640), Research Council of Norway “Toppforsk” Grant 249945, the Trond Mohn Foundation (Grant TMS2021TMT04), Marie Curie Grant MGATE (Grant 765549), and the National Infrastructure scheme of the Research Council of Norway - NORBRAIN (Grant 295721).

## AUTHOR CONTRIBUTIONS

L.A.L.D and C.K conceived the project; L.A.L.D obtained data with help from T.R and O.M.T.C; L.A.L.D and M.M.C performed the single-cell data analysis; W.P.C and L.A.L.D performed the population data analysis; W.P.C performed the decoder analysis; O.H developed the cell registration algorithm; M.J.S provided supervision to T.R and O.H; C.K provided supervision to L.A.L.D, M.M.C, O.M.T and acquired funding; L.A.L.D, W.P.C and C.K wrote the manuscript with inputs from all authors.

## COMPETING FINANCIAL INTERESTS

M.J.S was a scientific co-founder of Inscopix Inc, which produces the nVista miniature microscope used to acquire our neural calcium imaging data. This financial interest was fully removed partway through our study. The rest of the authors declare no competing interests.

## Methods

### Experimental model and subject details

Experiments were performed in compliance with the Norwegian Animal Welfare Act and the European Convention for the protection of Vertebrate Animals used for Experimental and Other Scientific Purposes. All experiments were approved by the Norwegian Food Safety Authority (Mattilsynet; protocol IDs #17898 and #30005). Transgenic animals expressing the calcium indicator GCaMP6s under control of the CamKII promoter (CaMKII-tTA; tetO-GCaMP6s) were used. Adult females (N = 7) were between 3 and 6 months of age at the time of the experiment. They were housed individually after surgery, and kept on a 12-h light-dark schedule with food and water provided ad libitum. Health was monitored daily, and experiments were always conducted during the dark phase of the light-dark cycle.

### Implant surgery

To gain optical access to the anterior cingulate cortex (ACC), we used a prism/GRIN lens assembly (ProView Integrated Prism Lens, Lens Length 4.3mm, Numerical Aperture 0.36, Pitch 1/2, Inscopix). Briefly, the procedure was performed under isoflurane anaesthesia (5% induction, 0.5-2.5% maintenance) and analgesia (Temgesic, Marcain, Metacam). We made a square craniotomy over the ACC with the following corners coordinates relative to bregma: A/P 0.2mm, M/L 0.2mm; A/P 0.2mm, M/L 1.4mm; A/P 1.4mm, M/L 0.2mm; A/P 1.4mm, M/L 1.4mm. The dura was carefully removed, and a dissecting knife (Fine Science Tools) was used to cut the brain along the medial axis of the craniotomy (D/V −2mm) to facilitate the insertion of the ProView Integrated Prism Lens (D/V −1.8mm). Any exposed brain tissue was covered with Kwik-Sil (World Precision Instruments), then the implant was secured to the skull using dental cement (Super-Bond C&B, SunMedical) and the surface of the GRIN lens covered with Kwik-Sil.

### Baseplate installation

Mice were given a minimum of 4 weeks after the implant surgery before being subjected to the baseplate installation. Mice were anaesthetised with isoflurane (5% induction, 0.5-2.5% maintenance), the protective layer of Kwik-Sil over the GRIN lens was removed, and the surface of the lens cleaned. A baseplate (Inscopix) was fitted on the miniaturised fluorescence microscope (“miniscope”, nVista 2, Inscopix), which was secured to the stereotax using a holder (Inscopix). The surface of the objective lens of the miniscope was aligned parallel to the surface of the GRIN lens, and the miniscope carefully lowered until we could see blood vessels in the field of view (FOV) using the nVista imaging software. Since GCaMP dynamics in the ACC under anaesthesia are quite low, we used blood vessels and toe-pinch evoked fluorescence dynamics to select a FOV. Once selected, we raised the miniscope 40µm to compensate for the shrinkage of the adhesive used to secure the baseplate to the skull (Venus Diamond Flow, Kulzer). After securing the baseplate, we added a custom-made aluminium headbar to allow for head-fixation in subsequent imaging sessions.

### Behavioural apparatus

#### Arena

We used a circular arena (diameter: 50cm, height: 40cm) containing a proximal cue card and placed on either a linoleum flooring (N = 2 mice) or a transparent acrylic sheet (N = 5 mice). We divided the arena in four quadrants, and an object was placed in the middle of two quadrants. The identity of the quadrants containing the objects was the same throughout the experiment and across mice, as well as the objects. To ensure that the objects placement was the same from one session to the next, we guided positioning using a paper pattern. The arena walls, the floor, and the objects were wiped with 70% ethanol before and after every session.

#### Behavioural recording

For mice exploring the arena on a transparent acrylic sheet, our behavioural stream was acquired through two cameras, one placed overhead and one underneath the transparent acrylic sheet (The Imaging Source). Both cameras recorded at 60Hz. For mice exploring the arena on a linoleum flooring, the behavioural stream was acquired through a webcam placed overhead and recording at the same frame rate as the imaging recording.

#### Calcium-imaging recording

All recordings were performed with a miniaturised head-mounted fluorescence microscope (nVista 2, Inscopix). The frame rate was 20 Hz and the LED power and gain were adjusted to each animal, but the power never above 1 mW to minimise photobleaching. On days with one recording session (Habituations 3 to 8), recordings lasted 30 minutes. On days where we had two sessions back to back (Open Field 1 and 2, Habituation 1 and 2), we acquired data for the first 15 minutes of each session to minimise photobleaching. For consistency during analysis, we analysed the first 15 minutes of each recording.

### Data processing

#### Behavioural movies - Position data

The position of the mouse in the arena throughout the recording sessions was extracted using DeepLabCut ^62,63^. Specifically, we labelled 30 frames taken from 26 videos across 3 animals then 95% was used for training. We used a ResNet-50-based neural network with default parameters for 200000 training iterations. We found the test error was: 4.65 pixels, train: 2.71 pixels (image size was 720 by 960). We then used a p-cutoff of 0.6 to condition the X,Y coordinates for future analysis. This network was then used to analyse videos from similar experimental settings. We then downsampled the position data with a factor 3 to match the sampling of the imaging recording.

#### Behavioural movies - Behaviour labelling

Behaviours were manually labelled using a Jython-based, custom-developed graphical user interface ^64^. We labelled the data from Day 1 to Day 7 for the 5 mice where we had both the overhead and the bottom-up behavioural steams. We first spatially downsampled the behavioural movies by a factor 2, then annotated the following behaviours frame by frame: rearing against the wall of the arena, resting, and grooming. We then used a custom-made script (MATLAB) to synchronise the behavioural labels to the neural data (See **Methods**, “Synchronisation between behavioural and imaging recordings”).

#### Imaging movies

Imaging movies were recorded in raw format and were first decompressed into hd5f and spatially downsampled with a factor 2 to reduce computational processing times and boost signal-to-noise (Inscopix Image Decompressor software). Once in hdf5 format, movies were processed through Ciatah ^65^. Recordings were further spatially downsampled with a factor 2. We performed motion correction by registering all frames in an imaging session to a chosen reference frame using TurboReg ^66^, and converted each movie to relative changes in fluorescence (ΔF/F) using the following formula:

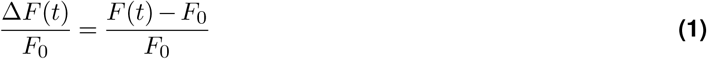

where F_0_ was the mean image over the entire movie.

#### Cell extraction and registration across sessions

We utilised TRACKER ^27^, an algorithm performing iterative cell extraction and registration, enabling us to track the same neurons throughout the experiment. Briefly, TRACKER is composed of a Cell Extraction module and a Cell Registration module, with a feedback mechanism between the two. Spatial footprints and fluorescence time series of putative cells are obtained from the Cell Extraction module - in our case by using EXTRACT ^67,68^. The Cell Registration module uses data extracted from the Cell Extraction module to align cells across sessions. We matched resulting putative cells across sessions based on Tanimoto similarity, defined as

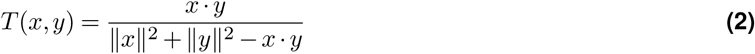

where x and y are vectors corresponding to flattened spatial filters. Next, a global cell repertoire (global cell map) is computed and the Cell Extraction module is reinitialised with the global cell map. This allowed EXTRACT to find cells that could have been missed on different sessions. We performed this procedure (cell extraction, across session registration and reinitialisation) for five iterations.

#### Synchronisation between behavioural and imaging recordings

For mice placed on the transparent acrylic sheet (N = 5 animals), behavioural and imaging streams were synchronised post hoc during the data processing. Briefly, we had placed LEDs within the field of view of the cameras, but outside of the arena explored by the mice, connected to an Arduino microcontroller generating 250ms pulses at random intervals. A logic analyzer (Saleae) was connected to the Arduino and the SYNC port of the nVista Digital Acquisition Box, recording the timing of the Arduino digital pulses and the imaging frames, in order to perform precise temporal alignments of the behavioural and imaging streams. A custom-made script (MATLAB) extracted the blinking LED pattern from the behavioural videos and matched it to the digital pulses pattern emitted by the Arduino and recorded by the logic analyzer, computing an “aligner” that was subsequently used to match imaging frames to behavioural frames. For mice placed on the linoleum flooring (N = 2 mice), the behavioural and imaging recordings were triggered to start simultaneously by ANY-maze, and recorded at the same frame rate.

### Activity maps

After cell extraction, we processed activity traces by keeping fluorescent events that were over 3 standard deviations of the mean fluorescence for a given cell and used Ciatah to extract peaks of activity from the filtered traces. Activity maps were generated by binning the location of each fluorescent event (2×2 cm bins) for each session, dividing the mean fluorescence in each bin by the time spent in that bin, and smoothing with a Gaussian.

### Object coding

We functionally characterised cells as object cells using a shuffling procedure. We shuffled the activity trace of the cells by circularly shifting the timestamps of the fluorescence trace, wrapping the end of the trace at the start, then generated an amplitude map. We repeated this 500 times per session per cell, then averaged the fluorescence rate in the 2×2 cm bins corresponding to the object locations and compared it to the real values. A cell was classified as an object cell for a given object if its true mean rate in the corresponding object bins was higher than a threshold set at the 95% percentile of the shuffled distribution. Using this method, we only classified cells as object cells if they were positively modulated by the object, and omitted negative modulation.

### Object score

Each cell was attributed an object score by applying a template matching procedure ^33^. Briefly, a Gaussian template was placed over the object’s location. We then calculated the Pearson correlation between the template and the real firing map of the cell. The object score gave us a metric of how much a cell’s firing field resembles the template field.

### Coarse graining

To look for correlated ensembles of neurons, we used a coarse-graining procedure, a method inspired by the renormalization group in statistical physics, where complex local interactions give rise to stable large-scale patterns. By applying coarse-graining, we filter out events at the microscopic scale (here, the activity of single cells) to reveal the emergence of large-scale, macroscopic patterns. At each iteration (K step), we computed the pairwise Pearson correlation matrix of all signals in the current pool. We then identified the most correlated pair, summed their activity traces, and normalized the resulting signal such that the mean of its nonzero values equaled one - preserving the meaning of zero (silence) while removing the amplitude increase introduced by summation ^35^. This pair was then removed from the pool and the process repeated until all signals had been paired. The normalized signals from one iteration served as the input to the next, so the procedure was applied recursively: at K step k, each coarse-grained signal comprised 2^(k-1)^ original neurons. Because this binary pairing requires an even number of inputs at each iteration, any residual unpaired signal at the end of a step was necessarily excluded from further iterations. These excluded cells, whilst anti-correlated to the rest of the population, could actually be quite informative because they diverge from the collective mean.

### Decoding

We used decision trees to decode either position-based or behaviour-based labels. We first built label curves (see Figure S7A, right) for each mouse and each session. The position labels were the following: edges of the left object, edges of the right object, centre of the left object, centre of the right object, edges of the arena. The behaviour labels were grooming, resting and rearing against the arena walls. Once we obtained the labels, we split the dataset in each session in training (80%) and testing (20%) data. We looped through each session and each coarse-graining step and grew a cross-validated decision tree with 10-fold cross-validation. For control, we performed a randomized permutation of the labels (“shuffled data”) before growing the tree. To obtain one value of error rate and maximum depth per session and clustering step, for both trees grew from real and shuffled data, we computed the median error rate and median maximum depth from the output of the 10-fold cross-validated trees. To identify the coarse-grained signals with the most and least weight in the decoder performance, we computed estimates of predictor importance on Day 1 at K8. For each animal and from the 10-fold cross-validated trees, we selected a unique coarse-grained signal as most weighted and a unique one as least weighted, using the most frequent signal that had the highest importance score. To obtain single-cells from these most and least weighted signals, we computed the correlations between the single-cells making up each signal, and the signal itself. Using this metric, we kept the single-cells that were most correlated, as we assumed they would be driving their respective assemblies. We used the 90th percentile as a threshold to select cells based on their correlation with the coarse-grained signal trace. This analysis can be performed on any day, but we specifically chose to look at the first day with objects in the environment since we could then compare the weight of single-cells to the coding properties they would exhibit in the remaining days of the experiment.

### Histological procedures and analysis

#### Sample preparation

At the end of the experiment, animals were transcardially perfused with 10X RNAse-free PBS and fixated with 4% RNAse-free PFA. The head was removed and immersed in PFA for 2 days with the implant still in place. Following fixation, the brain was dissected out and placed in a 30% RNAse-free sucrose solution for cryopreservation, then sectioned using a cryostat in several series of 30µm thick slices.

#### Native fluorescence

To assess the fluorescence signal from cells expressing GCaMP6s, we did not perform any signal enhancement and instead looked at the native fluorescence. We mounted a series of slices on glass slides with a mounting medium containing a DAPI stain (Fluoromount-G mounting medium, with DAPI, ThermoFisher Scientific).

## Supplementary Figures

**Figure S1.**
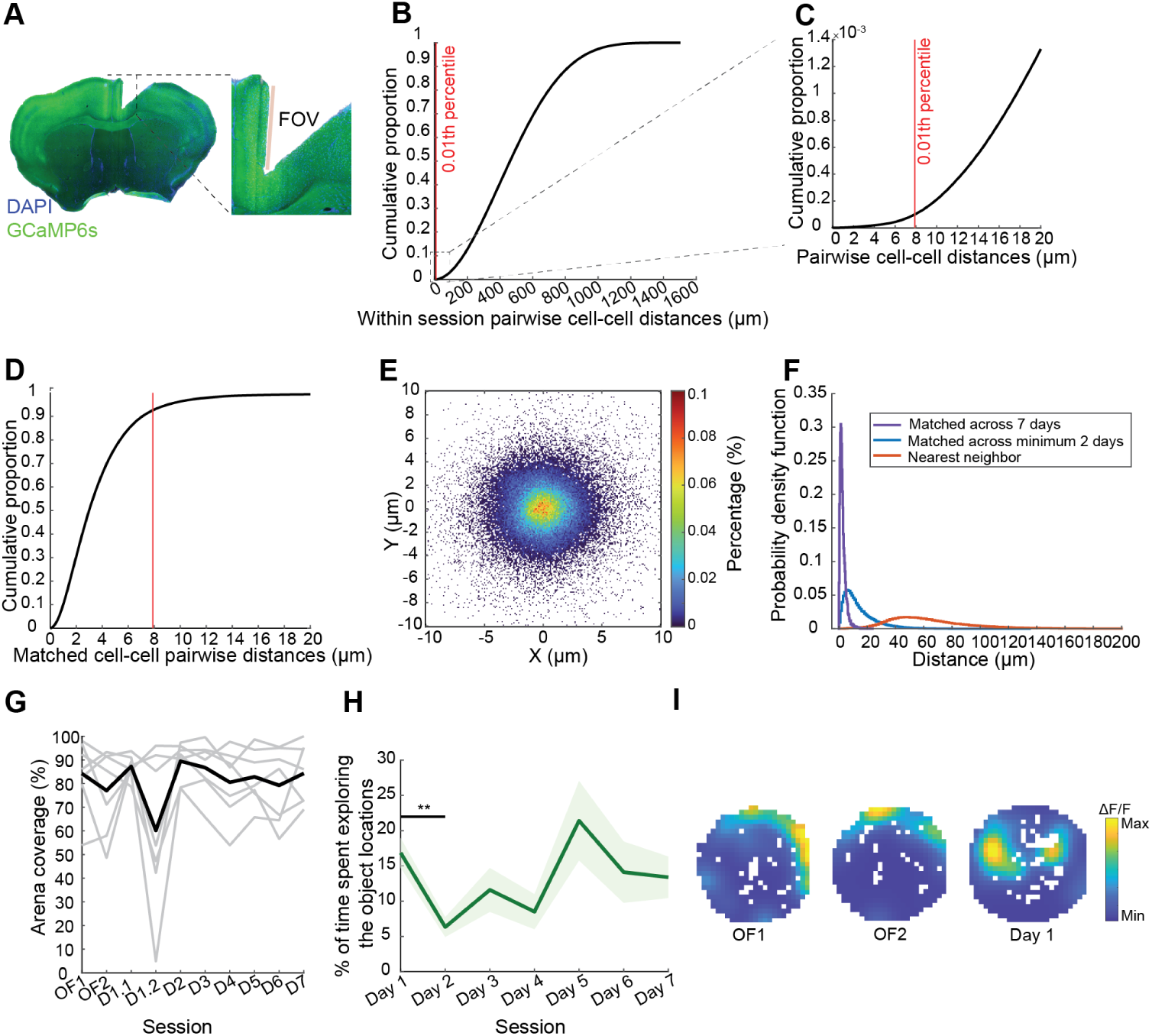
Optical access to ACC over repeated object exploration. **A)** Coronal brain slice (DAPI, blue; GCaMP6s, green) with tract resulting from the prism implant depicted in Figure 1A. **B)** Pairwise centroid Euclidean distances between cells, within a single session, for all imaging sessions across mice. Red line = 0.01^th^ percentile. **C)** Zoom from panel B) showing the cumulative proportion of neurons within 20µm of one another. **D)** Same metric as panel B) but restricted to neuron-neuron pairs grouped as the same cell across sessions under the matching procedure (see **Methods**). The red line is at the same location as in panels B) and C). **E)** For neurons matched with at least another session, displacement in X and Y (in pixels) from their average centroid location, represented as a percentage of occurrence. N = 73,049 individual cells pooled from 7 mice. **F)** Probability density function of the distance, in µm, between neurons matched across at least 2 sessions (blue), across all sessions of objects exploration (purple), and neurons and their nearest neighbors (orange). N = 134,834 distances between neurons matched across at least 2 sessions; N = 20,760 distance between neurons matches across all sessions of objects exploration; N = 162,592 distances between nearest neighbors. For statistical comparisons, we computed pairwise *d’* (discriminability index) of the log-transformed distances. *d’* matched across at least 2 sessions vs nearest neighbors = 2.45; *d’* matched across at least 2 sessions vs matched on all objects exploration sessions = 2.02; *d’* matched on all objects exploration sessions vs nearest neighbors = 5.01. **G)** Percentage of arena covered in each session, individual animals (gray) and averaged (black). **H)** Average time spent exploring the objects on each day, N = 7 mice. Data are represented as mean ± SEM (dark line = mean, shaded area = SEM). Day 1 vs Day 2: Two-sample t-test, *\*\*p* = 0.0044. **I)** Activity maps of the same cell in OF1, OF2, and Day 1. The cell did not exhibit any constrained activity in the open field sessions. However on Day 1, upon introduction of the objects, its activity was localized around the objects location and the cell was classified as a multi object-location cell. *\*\*p < 0.01*

**Figure S2.**
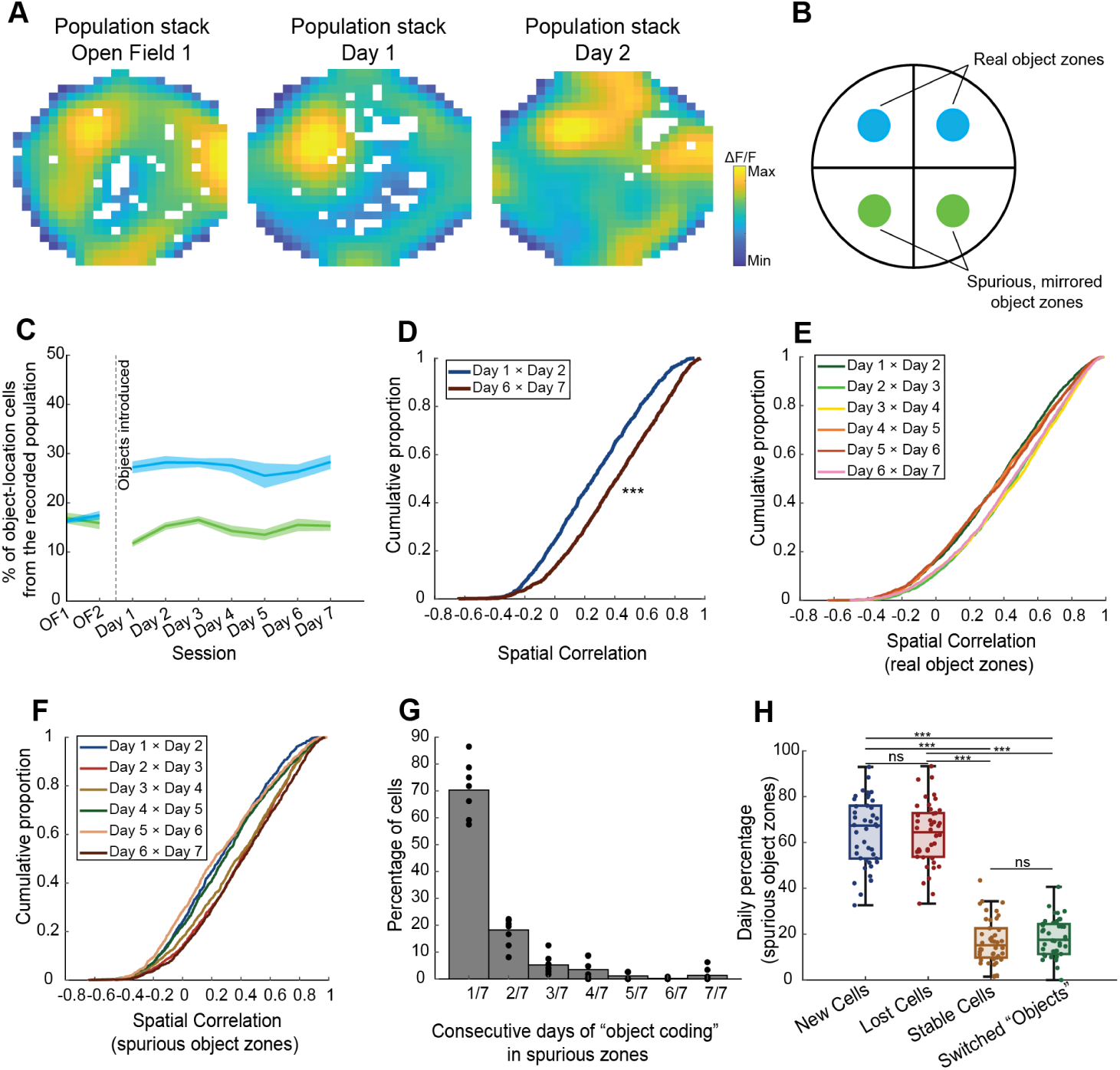
Using spurious object locations to disentangle object-location coding and spatial coding in ACC. **A)** Population stack of 705 cells registered across the first open field session and the first 2 days with objects in the environment, from one example animal. **B)** Schematics representation of the arena, the location of the objects, and the location of spurious object zones used as a control. **C)** Proportion of object-location cells in the spurious zones across Open Field sessions (OF1 and OF2 and throughout the remainder of the experiments once the objects have been introduced for real (blue) and spurious (green) object zones, N = 7 mice. Data are represented as mean ± SEM (dark line = mean, shaded area = SEM). For statistical comparisons of the spurious proportions between sessions, see Table S3. **D)** Spatial correlation score for object-location cells in the spurious object zones at the start (Day 1 × Day 2, dark blue) and the end of the familiarization to the objects (Day 6 × Day 7, burgundy). N = 7 mice. Two-sample Kolmogorov-Smirnov test, *\*\*\*p* = 3.9947e-13. **E)** Spatial correlation score for object-location cells in the real object zones, consecutive sessions comparisons throughout the experiment. **F)** Similar to E), but for spurious object zones. **G)** From the subset of unique spurious object-location cells registered throughout all object exploration sessions (n = 823 across 7 animals), percentage of object-location cells coding consecutively for the same spurious object locations, out of the 7 days of the experiment. Only 1.34% of cells coded for the same object throughout the 7 days, and 0.24% throughout 6 consecutive days. One dot = percentage for one animal. **H)** Using spurious object zones, percentages of new, lost, stable cells and cells switching their “object preference” in the spurious object-location cell ensemble between day X and day X+1. One dot = percentage between 2 consecutive sessions from Day 1 to Day 7 for a single animal, N = 7 mice. Pooled data are represented as median and interquartile range ± nonoutlier maximum/minimum. For statistical comparisons between groups, see Table S4. *\*\*\*p*<0.001

**Figure S3.**
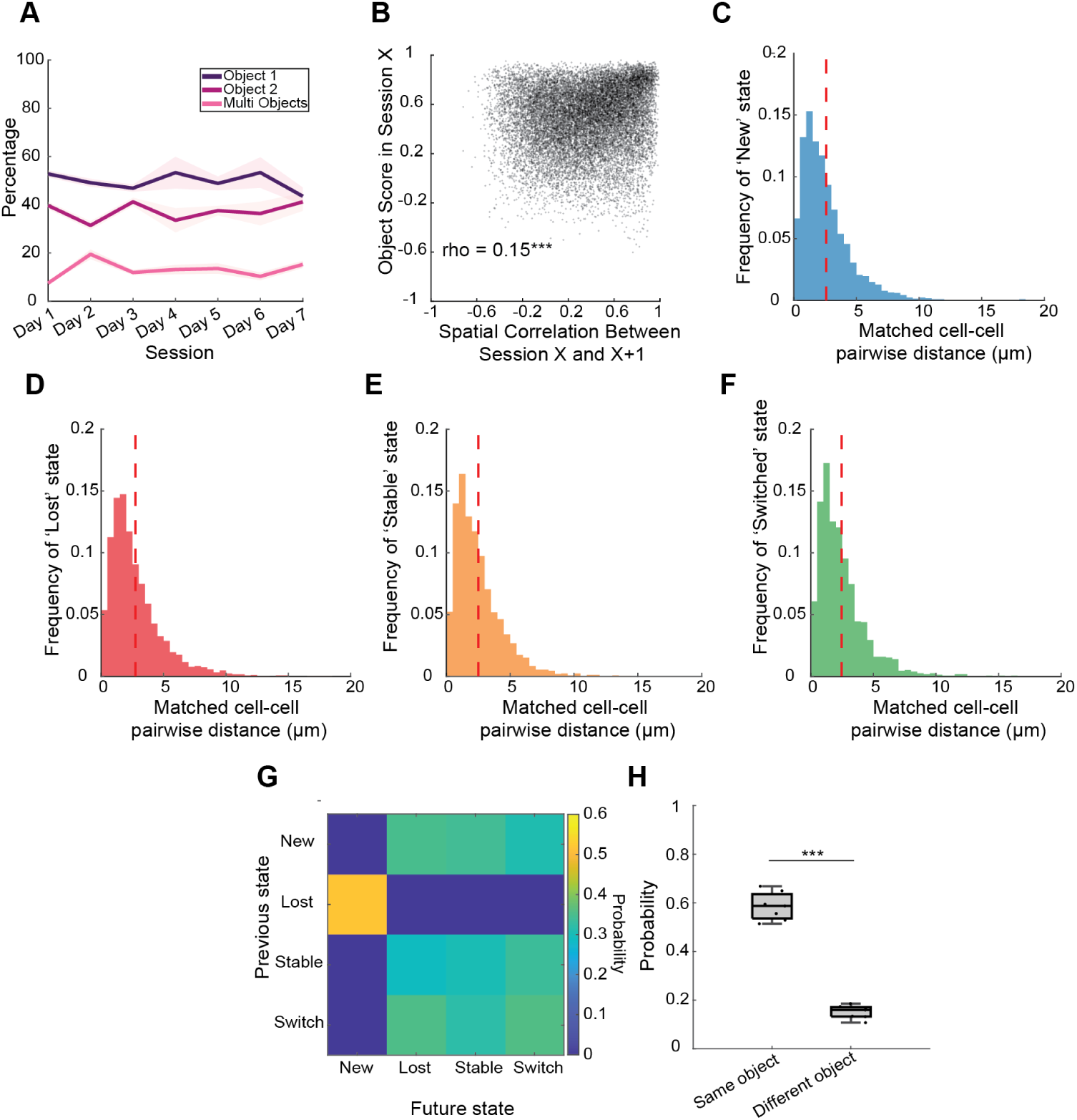
Single-cell dynamic coding is not an artefact of subpar cell matching across sessions. **A)** From the population of object-location cells, proportion of cells coding for object 1, object 2, and both objects (multi objects). N = 7 mice. Data are represented as mean ± SEM (dark line = mean, shaded area = SEM). **B)** Correlation between the object score in session X and the spatial correlation score between session X and X+1. One dot = one cell, N = 11477 cells. Spearman’s rho correlation, rho = 0.1500, *\*\*\*p* = 1.4592e-87. **C)** For cells qualifying as a “new” cell (see Figure 2D), distribution of the distance between the matched centroid locations between the 2 consecutive sessions where the cells qualified as “new” cells. Mean matched distance = 2.59µm, N = 2953 distances. **D)** Same as C for “lost” cells. Mean matched distance = 2.76µm, N = 2860 distances. **E)** Same as C for “stable” cells. Mean matched distance = 2.52µm, N = 2351 distances. **F)** Same as C for “switched” cells. Mean distance = 2.47µm, N = 1500 distances. In C, D, E and F the red bar denotes the mean distance. **G)** Uniform probabilities of transitioning between categories between day X and day X+1. **H)** For cells specifically leaving the object-location cells ensemble then joining it in a later session, probability to code for the same or a different object as the first time they qualified as an object cell in the experiment. One dot = probability for a single animal. Pooled data are represented as median and interquartile range ± maximum/minimum. Two-sample t-test, *\*\*\*p* = 5.0151e-10. *\*p* <0.05, *\*\*p* < 0.01, *\*\*\*p*<0.001

**Figure S4.**
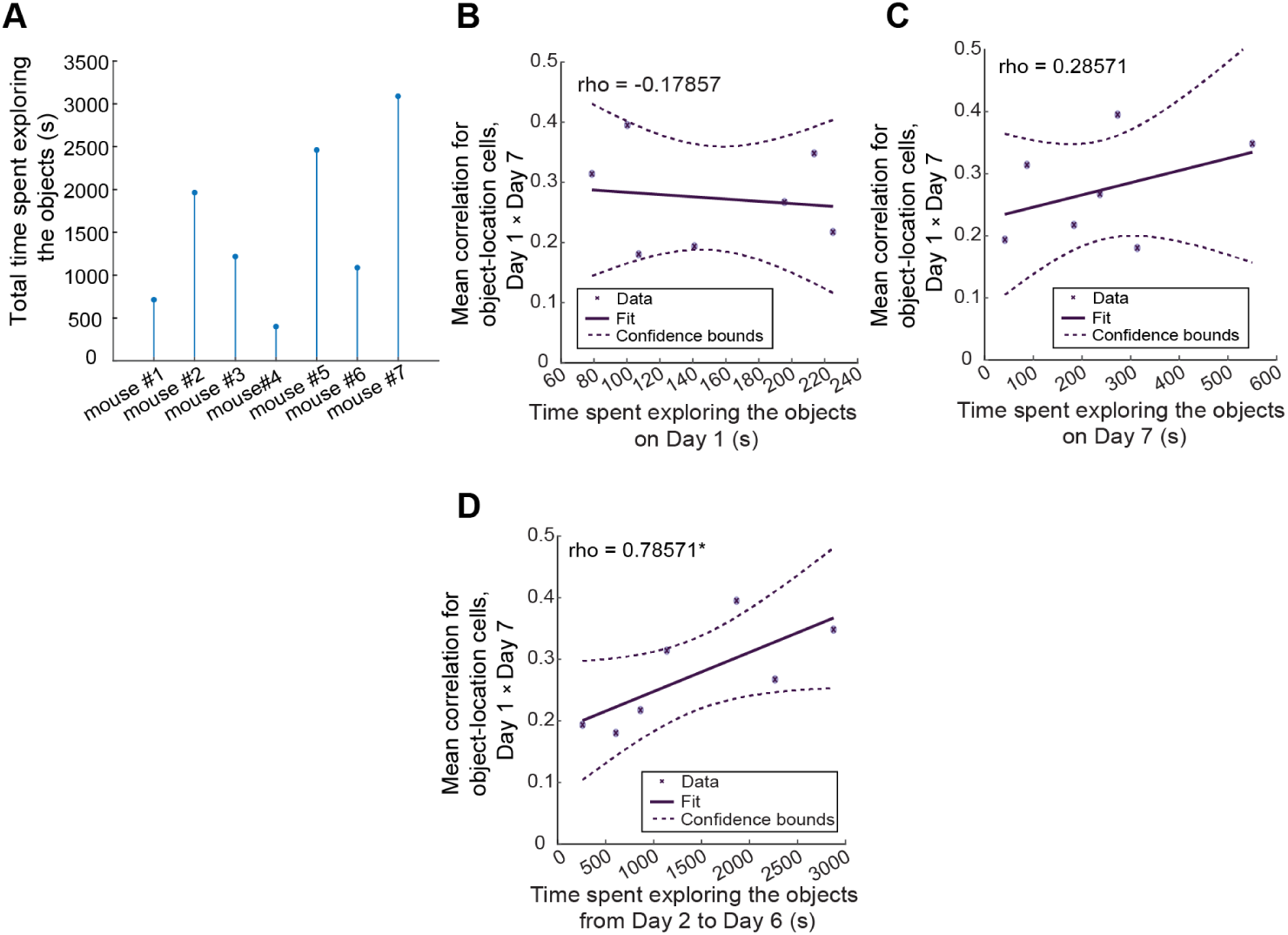
Single-cell coding stability is driven by experience and not mere sampling. **A)** Total time spent exploring the objects, in seconds, for each animal. **B)** Correlation between the mean spatial correlation score for object-location cells in Day 1 between their Day 1 and Day 7 fields, and the time the animal spent exploring the two objects on Day 1. Spearman’s rho correlation, rho = −0.1786, *p* = 0.7130. **C)** Correlation between the mean spatial correlation score for object-location cells in Day 1 between their Day 1 and Day 7 fields, and the time the animal spent exploring the two objects on Day 7. Spearman’s rho correlation, rho = 0.2857, *p* = 0.5559. **D)** Correlation between the mean spatial correlation score for object-location cells in Day 1 between their Day 1 and Day 7 fields, and the time the animal spent exploring the two objects from Day 2 to Day 6. Spearman’s rho correlation, rho = 0.78571, *\*p* = 0.0480. *\*p*<0.05

**Figure S5.**
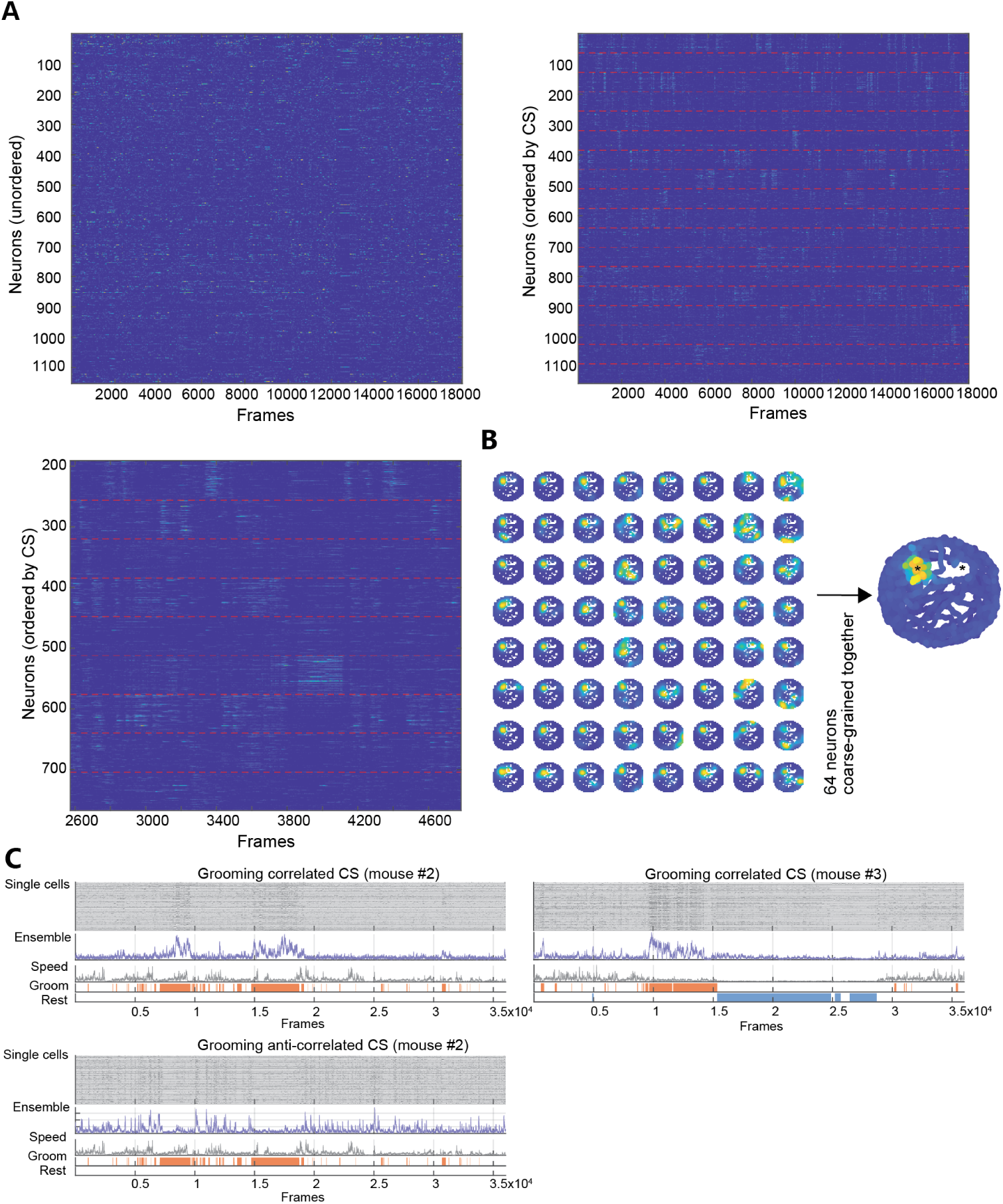
Population coding in ACC. **A)** Top left: Raster plot of activity during a recorded session (x axis), with no order to the neurons (y axis). Right: The same plot with the neurons reordered so they are grouped by coarse-grained signals (CS) i.e., cell ensembles, divided by red dotted lines. Bottom left: Zoom-in of the right panel. **B)** Activity maps of 64 neurons coarse-grained together at K7. **C)** Grooming-modulated activity in coarse-grained signals at K7. The top traces (grey, “single cells”) are from the 64 individual cells that have been coarse-grained together. The purple trace is the activity trace of the coarse-grain signal (“ensemble”), aligned to the speed of the animal and the scoring of two behavior variables (grooming, orange; resting, blue). Top and bottom left are assemblies included in Figure 3B, however because the animal did not perform any resting behavior in that session, the stationary position is a confound with grooming epochs. Right, example from a different animal that displayed both grooming and resting epochs in the same session. The ensemble is active when the mouse is grooming, but not resting, demonstrating that the activity is not driven by a stationary position or a low speed.

**Figure S6.**
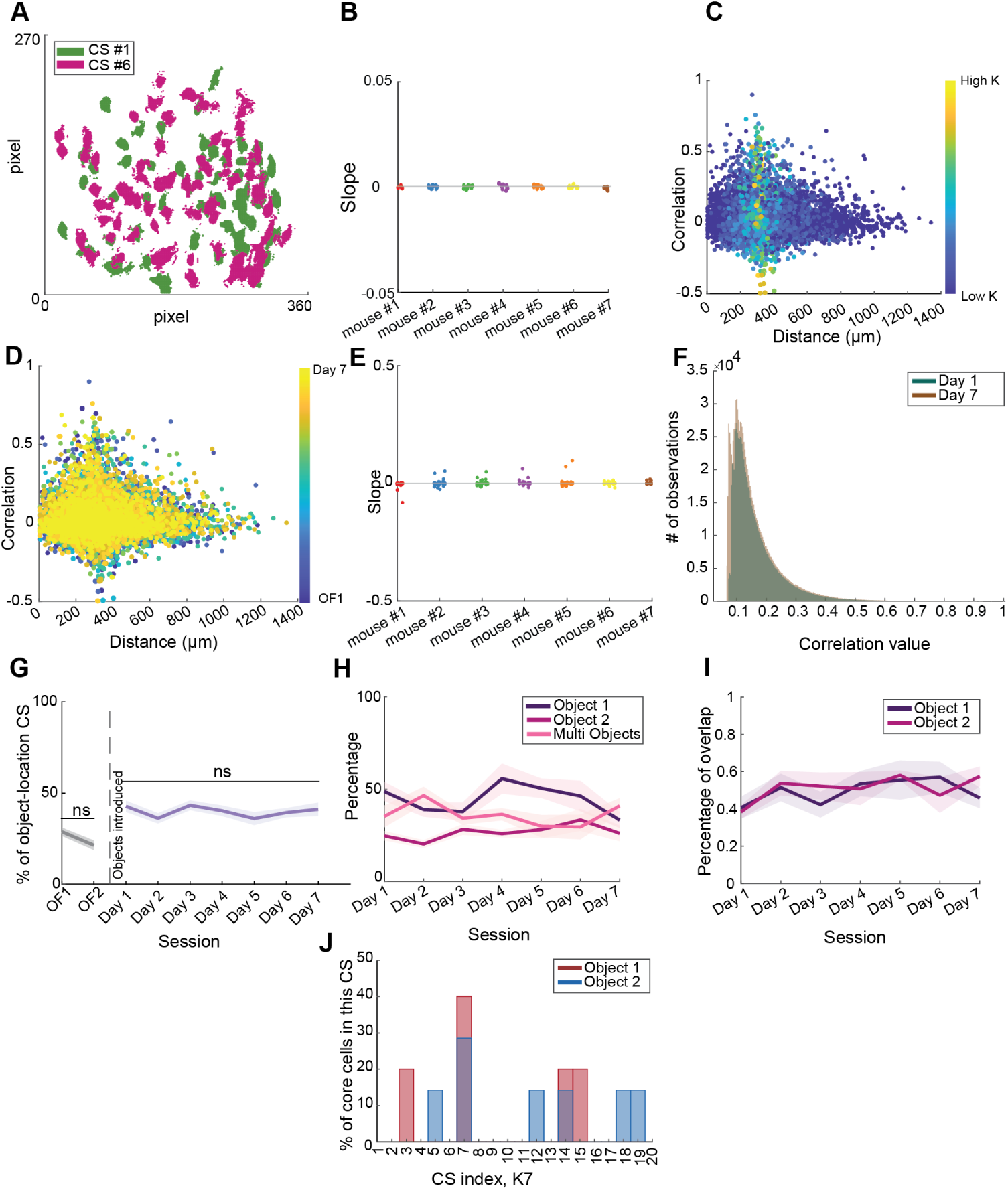
Population coding in ACC. **A)** Anatomical position in the FOV of neurons from two example coarse-grained signals. **B)** For each session and each mouse, we computed and plotted the slope of the correlation between 1) the anatomical distance between 2 neurons in the FOV and 2) their correlation value at step 7. **C)** Scatter plot of the correlation values between 2 neurons and their anatomical distance in the FOV, from low to high cluster steps (K), for an example mouse on a given day. **D)** Scatter plot of the correlation values between 2 neurons and their anatomical distance in the FOV, for all clustering steps and all sessions. For each session and each mouse, we computed the slope of the correlation between these 2 metrics and plotted it in **E)**. **F)** Top 10% correlation values between neurons on Day 1 (blue) and Day 7 (brown). Whilst the two distributions appear significantly different (Wilcoxon rank sum test, ranksum = 9.11e11, *\*\*\*p* <0.001), the effect size is very negligible (r = 0.069) due to the large sample size, suggesting that the distributions are practically equivalent. **G)** Proportion of object-location coding ensembles across the Open Field sessions (OF1 and OF2, grey, control sessions) and throughout the repeated familiarization to the objects (Day 1 to Day 7, purple), when using data that has been coarse-grained up to step 7, where ensembles are composed of 64 neurons. N = 7 mice. Dark line = mean, shaded area = SEM. For statistical comparisons between sessions, see Table S5. **H)** From the population of object-location coding ensembles, proportion of ensembles coding for object 1, object 2, and both objects (multi objects). N = 7 mice. Dark lines = mean, shaded areas = SEM. **I)** For cells coding constantly for the same object location throughout the experiment (extracted from Figure 2), we plotted the percentage of cells coarse-grained in the same assemblies at step 7. Mean object 1 core cells: 0.4951; mean object 2 core cells: 0.5119; mean all core cells combined: 0.5035). **J)** Representative example for a single mouse and a single session of the distribution of core cells throughout assemblies at step 7. Some assemblies contain core cells for both object 1 and object 2 (see CS #7 and CS #14). In G-I, data are represented as mean ± SEM (dark line = mean, shaded area = SEM)

**Figure S7.**
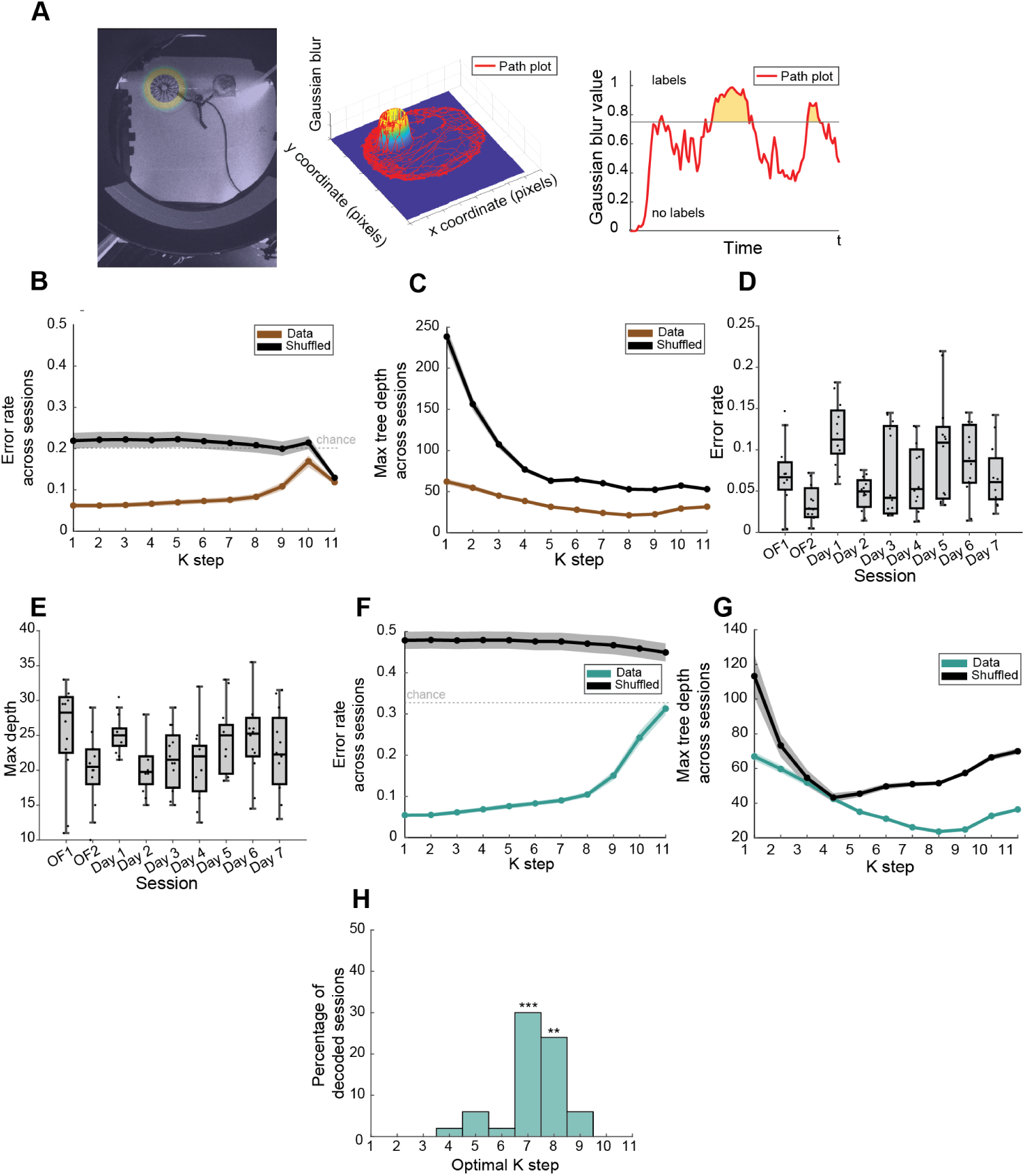
The optimal ensemble size is a feature of ACC and not specific to position-based, objects elicited coding. **A)** Schematics illustrating how the different zones (edges of right object, edges of left object, center of right object, center of left object, edges of the arena) were delimited, using the edges of the left object as an example. Left: a mask of the region of interest (ROI) is overlaid onto a frame from the camera recording the behavior of the animal. Middle: The path plot of the animal is overlaid on the ROI, represented as a 3D Gaussian blur. Right: When the path plot is over 75% of the Gaussian blur, it is used as a position label. **B)** Error rate at different clustering steps (K step) averaged across sessions when using decision trees to decode position-based labels. N = 7 mice. Real data brown, shuffled data black. Dark line = mean, shaded area = SEM. For statistical comparisons between real and shuffled data at each K step, see Table S6. For statistical comparisons between each K step for real data, see Table S7. **C)** Maximum tree depth at different clustering steps (K step) averaged across sessions when using decision trees to decode position-based labels. N = 7 mice. Real data brown, shuffled data black. Dark line = mean, shaded area = SEM. For statistical comparisons between real and shuffled data at each K step, see Table S8. For statistical comparisons between each K step for real data, see TableS9. **D)** Error rate at K7 and K8 when decoding position labels in all sessions. One dot = value of error rate for a single animal at either K7 or K8, as such there are 2 dots per animal for a given session. N = 7 mice. For statistical comparisons between sessions, see Table S10. **E)** Maximum depth at K7 and K8 when decoding position labels in all sessions. One dot = value of maximum depth for a single animal at either K7 or K8, as such there are 2 dots per animal for a given session. N = 7 mice. For statistical comparisons between sessions, see Table S11. **F)** Error rate at different clustering steps (K step) averaged across sessions when using decision trees to decode behavior-based labels. N = 5 mice. Real data blue, shuffled data black. Dark line = mean, shaded area = SEM. For statistical comparisons between real and shuffled data at each K step, see Table S12. For statistical comparisons between each K step for real data, see Table S13. **G)** Maximum tree depth at different clustering steps (K step) averaged across sessions when using decision trees to decode behavior-based labels. N = 5 mice. Real data blue, shuffled data black. Dark line = mean, shaded area = SEM. For statistical comparisons between real and shuffled data at each K step, see Table S14. For statistical comparisons between each K step for real data, see Table S15. **H)** Optimal ensemble size for decoding behavior data using decision trees. N = 5 animals. The decision trees were most efficient and accurate when using population data from K7 (64 neurons per ensemble, z-test, z-statistic = 4.1576, *\*\*\*p* = 3.216e-05) and K8 (128 neurons per ensemble, z-test, z-statistic = 2.7969, *\*\*p* = 5.16e-03). In B-C and F-G, data are represented as mean ± SEM (dark line = mean, shaded area = SEM). In D-E, pooled data are represented as median and interquartile range ± nonoutlier maximum/minimum. *\*\*\*p*<0.001

## Supplementary Tables

**Table S1.** ANOVA results for statistical comparisons between the proportion of object cells on each session. *\*p*<0.05, *\*\*p*<0.01, *\*\*\*p*<0.001.

| Group A | Group B | Lower Limit | A-B | Upper Limit | P-value |
| --- | --- | --- | --- | --- | --- |
| OF1 | OF2 | -9,2193 | -1,2148 | 6,7898 | 0,9999 |
| OF1 | Day 1 | -18,9698 | -10,9653 | -2,9607 | 0,0015** |
| OF1 | Day 2 | -19,9615 | -11,9569 | -3,9524 | 0,0004*** |
| OF1 | Day 3 | -19,9109 | -11,9064 | -3,9018 | 0,0004*** |
| OF1 | Day 4 | -19,3592 | -11,3547 | -3,3502 | 0,0009*** |
| OF1 | Day 5 | -17,276 | -9,2714 | -1,2669 | 0,0122* |
| OF1 | Day 6 | -18,0653 | -10,0608 | -2,0563 | 0,0047** |
| OF1 | Day 7 | -20,0613 | -12,0568 | -4,0523 | 0,0003*** |
| OF2 | Day 1 | -17,755 | -9,7505 | -1,746 | 0,0069** |
| OF2 | Day 2 | -18,7467 | -10,7422 | -2,7377 | 0,0019** |
| OF2 | Day 3 | -18,6961 | -10,6916 | -2,6871 | 0,0021** |
| OF2 | Day 4 | -18,1445 | -10,14 | -2,1355 | 0,0042** |
| OF2 | Day 5 | -16,0612 | -8,0567 | -0,0522 | 0,0474* |
| OF2 | Day 6 | -16,8506 | -8,8461 | -0,8415 | 0,02* |
| OF2 | Day 7 | -18,8465 | -10,842 | -2,8375 | 0,0017** |
| Day 1 | Day 2 | -8,9962 | -0,9917 | 7,0128 | 1 |
| Day 1 | Day 3 | -8,9456 | -0,9411 | 7,0634 | 1 |
| Day 1 | Day 4 | -8,394 | -0,3895 | 7,615 | 1 |
| Day 1 | Day 5 | -6,3107 | 1,6938 | 9,6983 | 0,9988 |
| Day 1 | Day 6 | -7,1001 | 0,9044 | 8,909 | 1 |
| Day 1 | Day 7 | -9,096 | -1,0915 | 6,913 | 1 |
| Day 2 | Day 3 | -7,9539 | 0,0506 | 8,0551 | 1 |
| Day 2 | Day 4 | -7,4023 | 0,6022 | 8,6067 | 1 |
| Day 2 | Day 5 | -5,319 | 2,6855 | 10,69 | 0,9742 |
| Day 2 | Day 6 | -6,1084 | 1,8961 | 9,9006 | 0,9974 |
| Day 2 | Day 7 | -8,1043 | -0,0998 | 7,9047 | 1 |
| Day 3 | Day 4 | -7,4529 | 0,5516 | 8,5561 | 1 |
| Day 3 | Day 5 | -5,3696 | 2,6349 | 10,6394 | 0,977 |
| Day 3 | Day 6 | -6,159 | 1,8455 | 9,8501 | 0,9978 |
| Day 3 | Day 7 | -8,1549 | -0,1504 | 7,8541 | 1 |
| Day 4 | Day 5 | -5,9212 | 2,0833 | 10,0878 | 0,995 |
| Day 4 | Day 6 | -6,7106 | 1,2939 | 9,2984 | 0,9998 |
| Day 4 | Day 7 | -8,7065 | -0,702 | 7,3025 | 1 |
| Day 5 | Day 6 | -8,7939 | -0,7894 | 7,2151 | 1 |
| Day 5 | Day 7 | -10,7898 | -2,7853 | 5,2192 | 0,9679 |
| Day 6 | Day 7 | -10,0005 | -1,996 | 6,0085 | 0,9962 |

**Table S2.** ANOVA results for statistical comparisons between proportions in each group forming the object cell ensemble. *\*p*<0.05, *\*\*p*<0.01, *\*\*\*p*<0.001.

| Group A | Group B | Lower Limit | A-B | Upper Limit | P-value |
| --- | --- | --- | --- | --- | --- |
| New | Lost | -5,2107 | 0,7253 | 6,6613 | 0,9890 |
| New | Stable | 3,7199 | 9,6559 | 15,5919 | 1,72E-04 *** |
| New | Switched | 15,5988 | 21,5348 | 27,4708 | 1,11E-20 *** |
| Lost | Stable | 2,9946 | 8,9306 | 14,8666 | 6,42E-04 *** |
| Lost | Switched | 14,8735 | 20,8095 | 26,7455 | 3,29E-19 *** |
| Stable | Switched | 5,9429 | 11,8789 | 17,8149 | 1,63E-06 *** |

**Table S3.**
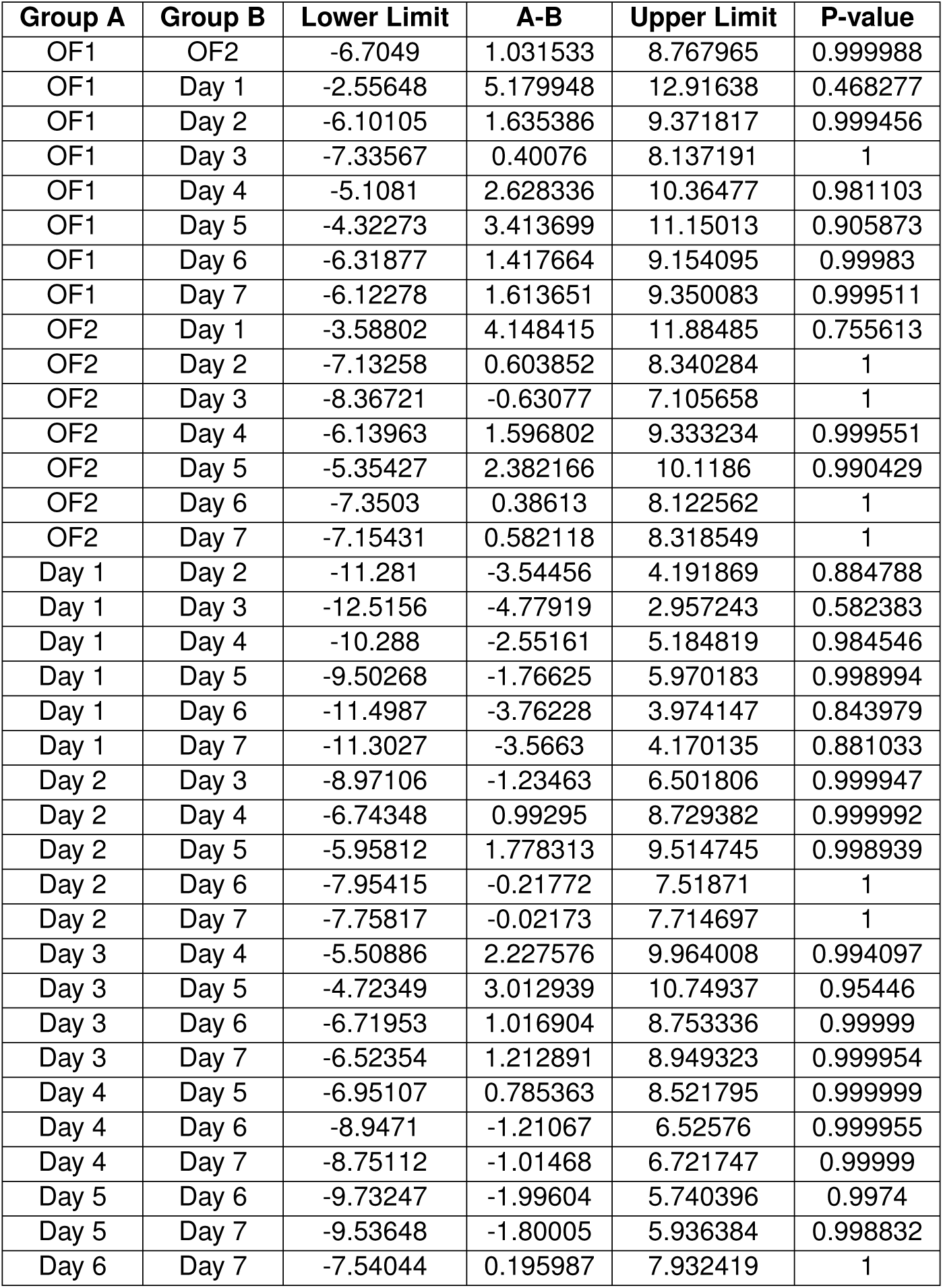
ANOVA results for statistical comparisons between the proportion of object-location cells, using spurious object locations, on each session.

**Table S4.** ANOVA results for statistical comparisons between proportions in each group forming the object-location cell ensemble, using spurious object locations. *\*p*<0.05, *\*\*p*<0.01, *\*\*\*p*<0.001.

| Group A | Group B | Lower Limit | A-B | Upper Limit | P-value |
| --- | --- | --- | --- | --- | --- |
| New | Lost | -5,6454 | 1.0262 | 7.6977 | 0.9791 |
| New | Stable | 41.0288 | 47.7003 | 54.3719 | <0.001*** |
| New | Switched | 39.4869 | 46.1585 | 52.8300 | <0.001*** |
| Lost | Stable | 40.0027 | 46.6742 | 53.3457 | <0.001*** |
| Lost | Switched | 38.4608 | 45.1323 | 51.80038 | <0.001*** |
| Stable | Switched | -8.2134 | -1.5419 | 5.1296 | 0.9340 |

**Table S5.** ANOVA results for statistical comparisons between the proportion of object cells on each session when using coarse-grained data from step 7 (ensemble size = 64 neurons). *\*p*<0.05, *\*\*p*<0.01.

| Group A | Group B | Lower Limit | A-B | Upper Limit | P-value |
| --- | --- | --- | --- | --- | --- |
| OF1 | OF2 | -8,7533 | 7,4245 | 23,6023 | 0,8585 |
| OF1 | Day 1 | -30,1843 | -14,0065 | 2,1712 | 0,1402 |
| OF1 | Day 2 | -23,5586 | -7,3808 | 8,7969 | 0,8624 |
| OF1 | Day 3 | -30,7407 | -14,5629 | 1,6148 | 0,1094 |
| OF1 | Day 4 | -27,7356 | -11,5579 | 4,6199 | 0,3556 |
| OF1 | Day 5 | -23,4381 | -7,2603 | 8,9174 | 0,8728 |
| OF1 | Day 6 | -26,7016 | -10,5238 | 5,6540 | 0,4830 |
| OF1 | Day 7 | -28,5499 | -12,3722 | 3,8056 | 0,2690 |
| OF2 | Day 1 | -37,6088 | -21,4311 | -5,2533 | 0,0023 |
| OF2 | Day 2 | -30,9831 | -14,8053 | 1,3724 | 0,0979 * |
| OF2 | Day 3 | -38,1652 | -21,9874 | -5,8097 | 0,0016 ** |
| OF2 | Day 4 | -35,1601 | -18,9824 | -2,8046 | 0,0106 * |
| OF2 | Day 5 | -30,8626 | -14,6849 | 1,4929 | 0,1035 |
| OF2 | Day 6 | -34,1261 | -17,9483 | -1,7705 | 0,0192 * |
| OF2 | Day 7 | -35,9744 | -19,7967 | -3,6189 | 0,0065 ** |
| Day 1 | Day 2 | -9,5521 | 6,6257 | 22,8035 | 0,9199 |
| Day 1 | Day 3 | -16,7342 | -0,5564 | 15,6214 | 1 |
| Day 1 | Day 4 | -13,7291 | 2,4487 | 18,6264 | 0,9999 |
| Day 1 | Day 5 | -9,4316 | 6,7462 | 22,9240 | 0,9120 |
| Day 1 | Day 6 | -12,6950 | 3,4827 | 19,6605 | 0,9987 |
| Day 1 | Day 7 | -14,5434 | 1,6344 | 17,8122 | 1 |
| Day 2 | Day 3 | -23,3599 | -7,1821 | 8,9957 | 0,8794 |
| Day 2 | Day 4 | -20,3548 | -4,1770 | 12,0007 | 0,9952 |
| Day 2 | Day 5 | -16,0573 | 0,1205 | 16,2983 | 1 |
| Day 2 | Day 6 | -19,3207 | -3,1430 | 13,0348 | 0,9994 |
| Day 2 | Day 7 | -21,1691 | -4,9913 | 11,1864 | 0,9846 |
| Day 3 | Day 4 | -13,1727 | 3,0051 | 19,1828 | 0,9995 |
| Day 3 | Day 5 | -8,8752 | 7,3026 | 23,4804 | 0,8692 |
| Day 3 | Day 6 | -12,1386 | 4,0391 | 20,2169 | 0,9962 |
| Day 3 | Day 7 | -13,9870 | 2,1908 | 18,3685 | 1 |
| Day 4 | Day 5 | -11,8803 | 4,2975 | 20,4753 | 0,9942 |
| Day 4 | Day 6 | -15,1437 | 1,0341 | 17,2118 | 1 |
| Day 4 | Day 7 | -16,9921 | -0,8143 | 15,3635 | 1 |
| Day 5 | Day 6 | -19,4412 | -3,2635 | 12,9143 | 0,9992 |
| Day 5 | Day 7 | -21,2896 | -5,1118 | 11,0660 | 0,9821 |
| Day 6 | Day 7 | -18,0261 | -1,8484 | 14,3294 | 1 |

**Table S6.**
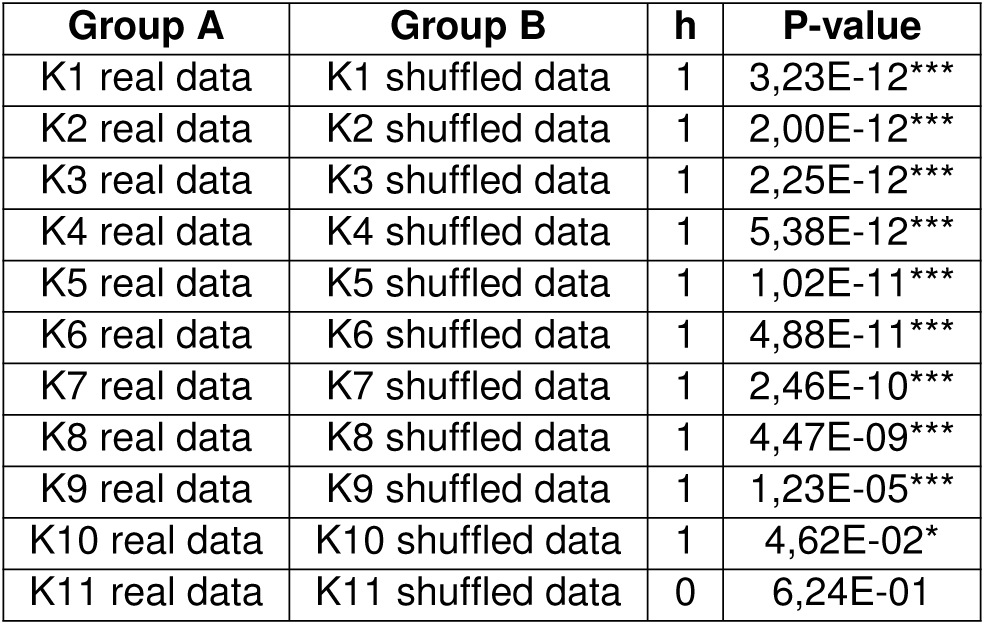
Two-sample t-test results for statistical comparisons at each K step of the error rate between real and shuffled data, position-based labels. *\*p*<0.05, *\*\*\*p*<0.001.

**Table S7.** Statistical comparisons between the error rates at each K step for real data, position-based labels. *\*p*<0.05, *\*\*\*p*<0.001.

| Group A | Group B | Lower Limit | A-B | Upper Limit | P-value |
| --- | --- | --- | --- | --- | --- |
| K1 | K2 | -0,0339 | 0,0000 | 0,0339 | 1 |
| K1 | K3 | -0,0352 | -0,0013 | 0,0326 | 1 |
| K1 | K4 | -0,0383 | -0,0044 | 0,0296 | 1 |
| K1 | K5 | -0,0417 | -0,0078 | 0,0261 | 0,9997 |
| K1 | K6 | -0,0447 | -0,0108 | 0,0231 | 0,9949 |
| K1 | K7 | -0,0476 | -0,0137 | 0,0203 | 0,9696 |
| K1 | K8 | -0,0551 | -0,0212 | 0,0128 | 0,6429 |
| K1 | K9 | -0,0803 | -0,0464 | -0,0125 | 0,0006*** |
| K1 | K10 | -0,1423 | -0,1070 | -0,0717 | 0,0000*** |
| K1 | K11 | -0,1243 | -0,0565 | 0,0114 | 0,2094 |
| K2 | K3 | -0,0352 | -0,0013 | 0,0326 | 1 |
| K2 | K4 | -0,0383 | -0,0044 | 0,0295 | 1 |
| K2 | K5 | -0,0417 | -0,0078 | 0,0261 | 0,9997 |
| K2 | K6 | -0,0447 | -0,0108 | 0,0231 | 0,9949 |
| K2 | K7 | -0,0476 | -0,0137 | 0,0202 | 0,9693 |
| K2 | K8 | -0,0551 | -0,0212 | 0,0127 | 0,6417 |
| K2 | K9 | -0,0803 | -0,0464 | -0,0125 | 0,0005*** |
| K2 | K10 | -0,1423 | -0,1070 | -0,0717 | 0,0000*** |
| K2 | K11 | -0,1243 | -0,0565 | 0,0114 | 0,2090 |
| K3 | K4 | -0,0370 | -0,0031 | 0,0308 | 1 |
| K3 | K5 | -0,0404 | -0,0065 | 0,0274 | 0,9999 |
| K3 | K6 | -0,0434 | -0,0095 | 0,0244 | 0,9982 |
| K3 | K7 | -0,0463 | -0,0124 | 0,0215 | 0,9851 |
| K3 | K8 | -0,0538 | -0,0199 | 0,0140 | 0,7259 |
| K3 | K9 | -0,0790 | -0,0451 | -0,0112 | 0,0009*** |
| K3 | K10 | -0,1410 | -0,1057 | -0,0704 | 0,0000*** |
| K3 | K11 | -0,1230 | -0,0552 | 0,0127 | 0,2391 |
| K4 | K5 | -0,0374 | -0,0034 | 0,0305 | 1 |
| K4 | K6 | -0,0403 | -0,0064 | 0,0275 | 0,9999 |
| K4 | K7 | -0,0432 | -0,0093 | 0,0246 | 0,9985 |
| K4 | K8 | -0,0507 | -0,0168 | 0,0171 | 0,8854 |
| K4 | K9 | -0,0760 | -0,0420 | -0,0081 | 0,0032** |
| K4 | K10 | -0,1379 | -0,1026 | -0,0673 | 0,0000*** |
| K4 | K11 | -0,1199 | -0,0521 | 0,0158 | 0,3214 |
| K5 | K6 | -0,0369 | -0,0030 | 0,0309 | 1 |
| K5 | K7 | -0,0398 | -0,0059 | 0,0281 | 1 |
| K5 | K8 | -0,0473 | -0,0134 | 0,0206 | 0,9740 |
| K5 | K9 | -0,0725 | -0,0386 | -0,0047 | 0,0113* |
| K5 | K10 | -0,1345 | -0,0992 | -0,0639 | 0,0000*** |
| K5 | K11 | -0,1165 | -0,0487 | 0,0192 | 0,4276 |
| K6 | K7 | -0,0368 | -0,0029 | 0,0310 | 1 |
| K6 | K8 | -0,0443 | -0,0104 | 0,0235 | 0,9963 |
| K6 | K9 | -0,0695 | -0,0356 | -0,0017 | 0,0302* |
| K6 | K10 | -0,1315 | -0,0962 | -0,0609 | 0,0000*** |
| K6 | K11 | -0,1135 | -0,0457 | 0,0222 | 0,5287 |
| K7 | K8 | -0,0414 | -0,0075 | 0,0264 | 0,9998 |
| K7 | K9 | -0,0667 | -0,0327 | 0,0012 | 0,0700 |
| K7 | K10 | -0,1286 | -0,0933 | -0,0580 | 0,0000*** |
| K7 | K11 | -0,1107 | -0,0428 | 0,0251 | 0,6275 |
| K8 | K9 | -0,0592 | -0,0252 | 0,0087 | 0,3706 |
| K8 | K10 | -0,1211 | -0,0858 | -0,0505 | 0,0000*** |
| K8 | K11 | -0,1031 | -0,0353 | 0,0326 | 0,8488 |
| K9 | K10 | -0,0959 | -0,0606 | -0,0253 | 0,0000*** |
| K9 | K11 | -0,0779 | -0,0101 | 0,0578 | 1 |
| K10 | K11 | -0,0180 | 0,0505 | 0,1191 | 0,3849 |

**Table S8.**
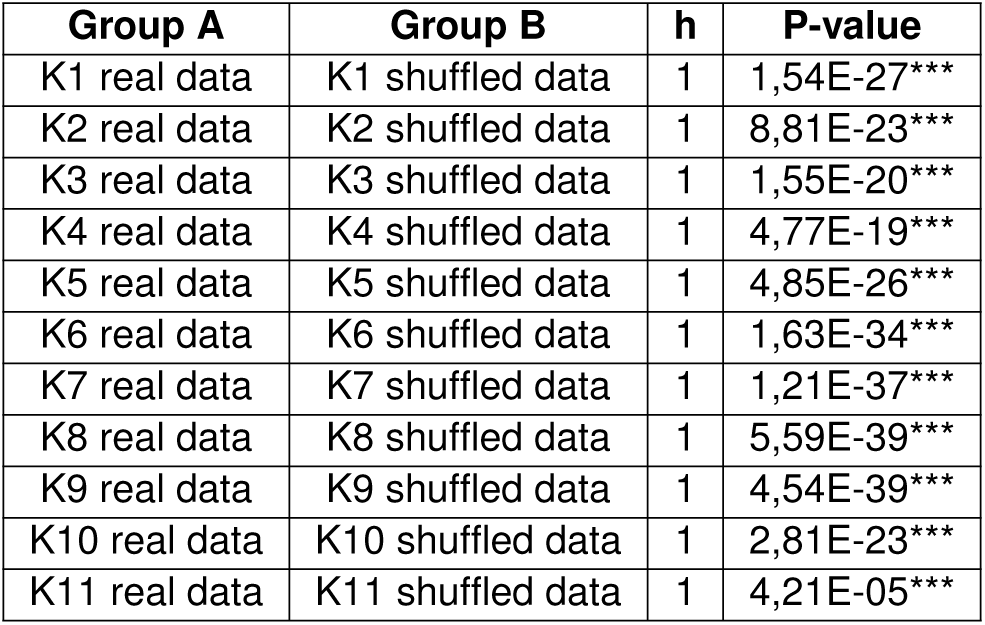
Two-sample t-test results for statistical comparisons at each K step of the maximum tree depth between real and shuffled data, position-based labels. *\*\*\*p*<0.001.

**Table S9.** Statistical comparisons between the maximum tree depths at each K step for real data, position-based labels; *\*p*<0.05, *\*\*p*<0.01, *\*\*\*p*<0.001.

| Group A | Group B | Lower Limit | A-B | Upper Limit | P-value |
| --- | --- | --- | --- | --- | --- |
| K1 | K2 | -0,8192 | 7,5204 | 15,8600 | 1,22E-01 |
| K1 | K3 | 8,6808 | 17,0204 | 25,3600 | 2,40E-09 |
| K1 | K4 | 15,2931 | 23,6327 | 31,9722 | 2,97E-19 |
| K1 | K5 | 22,3135 | 30,6531 | 38,9926 | 0,00E+00 |
| K1 | K6 | 25,7727 | 34,1122 | 42,4518 | 0,00E+00 |
| K1 | K7 | 29,7625 | 38,1020 | 46,4416 | 0,00E+00 |
| K1 | K8 | 32,4155 | 40,7551 | 49,0947 | 0,00E+00 |
| K1 | K9 | 31,2421 | 39,5816 | 47,9212 | 0,00E+00 |
| K1 | K10 | 23,9832 | 32,6633 | 41,3434 | 0,00E+00 |
| K1 | K11 | 13,8413 | 30,5204 | 47,1996 | 2,03E-07 |
| K2 | K3 | 1,1604 | 9,5 | 17,8396 | 1,11E-02 |
| K2 | K4 | 7,7727 | 16,1122 | 24,4518 | 2,54E-08 |
| K2 | K5 | 14,7931 | 23,1327 | 31,4722 | 2,53E-18 |
| K2 | K6 | 18,2523 | 26,5918 | 34,9314 | 0,00E+00 |
| K2 | K7 | 22,2421 | 30,5816 | 38,9212 | 0,00E+00 |
| K2 | K8 | 24,8951 | 33,2347 | 41,5743 | 0,00E+00 |
| K2 | K9 | 23,7217 | 32,0612 | 40,4008 | 0,00E+00 |
| K2 | K10 | 16,4628 | 25,1429 | 33,8230 | 2,78E-20 |
| K2 | K11 | 6,3209 | 23,0000 | 39,6791 | 4,68E-04 |
| K3 | K4 | -1,7273 | 6,6122 | 14,9518 | 2,74E-01 |
| K3 | K5 | 5,2931 | 13,6327 | 21,9722 | 7,68E-06 |
| K3 | K6 | 8,7523 | 17,0918 | 25,4314 | 1,98E-09 |
| K3 | K7 | 12,7421 | 21,0816 | 29,4212 | 7,28E-15 |
| K3 | K8 | 15,3951 | 23,7347 | 32,0743 | 1,93E-19 |
| K3 | K9 | 14,2217 | 22,5612 | 30,9008 | 2,62E-17 |
| K3 | K10 | 6,9628 | 15,6429 | 24,3230 | 3,50E-07 |
| K3 | K11 | -3,1791 | 13,5 | 30,1791 | 2,46E-01 |
| K4 | K5 | -1,3192 | 7,0204 | 15,3600 | 1,95E-01 |
| K4 | K6 | 2,1400 | 10,4796 | 18,8192 | 2,57E-03 |
| K4 | K7 | 6,1298 | 14,4694 | 22,8090 | 1,25E-06 |
| K4 | K8 | 8,7829 | 17,1224 | 25,4620 | 1,82E-09 |
| K4 | K9 | 7,6094 | 15,9490 | 24,2886 | 3,83E-08 |
| K4 | K10 | 0,3505 | 9,0306 | 17,7107 | 3,32E-02 |
| K4 | K11 | -9,7914 | 6,8878 | 23,5669 | 9,64E-01 |
| K5 | K6 | -4,8804 | 3,4592 | 11,7988 | 9,63E-01 |
| K5 | K7 | -0,8906 | 7,4490 | 15,7886 | 1,31E-01 |
| K5 | K8 | 1,7625 | 10,1020 | 18,4416 | 4,62E-03 |
| K5 | K9 | 0,5890 | 8,9286 | 17,2681 | 2,41E-02 |
| K5 | K10 | -6,6699 | 2,0102 | 10,6903 | 1,00E+00 |
| K5 | K11 | -16,8118 | -0,1327 | 16,5465 | 1,00E+00 |
| K6 | K7 | -4,3498 | 3,9898 | 12,3294 | 9,06E-01 |
| K6 | K8 | -1,6967 | 6,6429 | 14,9824 | 2,68E-01 |
| K6 | K9 | -2,8702 | 5,4694 | 13,8090 | 5,69E-01 |
| K6 | K10 | -10,1291 | -1,4490 | 7,2311 | 1,00E+00 |
| K6 | K11 | -20,2710 | -3,5918 | 13,0873 | 1,00E+00 |
| K7 | K8 | -5,6865 | 2,6531 | 10,9926 | 9,95E-01 |
| K7 | K9 | -6,8600 | 1,4796 | 9,8192 | 1,00E+00 |
| K7 | K10 | -14,1189 | -5,4388 | 3,2413 | 6,37E-01 |
| K7 | K11 | -24,2608 | -7,5816 | 9,0975 | 9,32E-01 |
| K8 | K9 | -9,5130 | -1,1735 | 7,1661 | 1,00E+00 |
| K8 | K10 | -16,7719 | -8,0918 | 0,5883 | 9,40E-02 |
| K8 | K11 | -26,9138 | -10,2347 | 6,4444 | 6,67E-01 |
| K9 | K10 | -15,5985 | -6,9184 | 1,7617 | 2,67E-01 |
| K9 | K11 | -25,7404 | -9,0612 | 7,6179 | 8,10E-01 |
| K10 | K11 | -18,9948 | -2,1429 | 14,7091 | 1,00E+00 |

**Table S10.** ANOVA results for statistical comparisons between sessions of the error rate at K7 and K8. *\*p*<0.05, *\*\*p*<0.01, *\*\*\*p*<0.001.

| Group A | Group B | Lower Limit | A-B | Upper Limit | P-value |
| --- | --- | --- | --- | --- | --- |
| OF1 | OF2 | -0.0155 | 0.0342 | 0.0839 | 0.4292 |
| OF1 | Day 1 | -0.1001 | -0.0503 | -0.0006 | 0.0448* |
| OF1 | Day 2 | -0.0290 | 0.0207 | 0.0704 | 0.9246 |
| OF1 | Day 3 | -0.0545 | -0.0047 | 0.0450 | 1.0000 |
| OF1 | Day 4 | -0.0467 | 0.0031 | 0.0528 | 1.0000 |
| OF1 | Day 5 | -0.0808 | -0.0311 | 0.0187 | 0.5646 |
| OF1 | Day 6 | -0.0681 | -0.0183 | 0.0314 | 0.9622 |
| OF1 | Day 7 | -0.0482 | 0.0015 | 0.0513 | 1.0000 |
| OF2 | Day 1 | -0.1343 | -0.0846 | -0.0348 | <0.001*** |
| OF2 | Day 2 | -0.0632 | -0.0135 | 0.0362 | 0.9946 |
| OF2 | Day 3 | -0.0887 | -0.0389 | 0.0108 | 0.2549 |
| OF2 | Day 4 | -0.0809 | -0.0311 | 0.0186 | 0.5611 |
| OF2 | Day 5 | -0.1150 | -0.0653 | -0.0155 | 0.0020** |
| OF2 | Day 6 | -0.1023 | -0.0525 | -0.0028 | 0.0300* |
| OF2 | Day 7 | -0.0824 | -0.0327 | 0.0171 | 0.4941 |
| Day 1 | Day 2 | 0.0213 | 0.0711 | 0.1208 | 0.0005*** |
| Day 1 | Day 3 | -0.0041 | 0.0456 | 0.0953 | 0.0996 |
| Day 1 | Day 4 | 0.0037 | 0.0534 | 0.1032 | 0.0253* |
| Day 1 | Day 5 | -0.0304 | 0.0193 | 0.0690 | 0.9491 |
| Day 1 | Day 6 | -0.0177 | 0.0320 | 0.0818 | 0.5223 |
| Day 1 | Day 7 | 0.0022 | 0.0519 | 0.1016 | 0.0339* |
| Day 2 | Day 3 | -0.0752 | -0.0254 | 0.0243 | 0.7936 |
| Day 2 | Day 4 | -0.0674 | -0.0176 | 0.0321 | 0.9700 |
| Day 2 | Day 5 | -0.1015 | -0.0518 | -0.0020 | 0.0346* |
| Day 2 | Day 6 | -0.0888 | -0.0390 | 0.0107 | 0.2523 |
| Day 2 | Day 7 | -0.0689 | -0.0192 | 0.0306 | 0.9509 |
| Day 3 | Day 4 | -0.0419 | 0.0078 | 0.0575 | 0.9999 |
| Day 3 | Day 5 | -0.0760 | -0.0263 | 0.0234 | 0.7618 |
| Day 3 | Day 6 | -0.0633 | -0.0136 | 0.0361 | 0.9944 |
| Day 3 | Day 7 | -0.0435 | 0.0063 | 0.0560 | 1.0000 |
| Day 4 | Day 5 | -0.0839 | -0.0341 | 0.0156 | 0.4326 |
| Day 4 | Day 6 | -0.0711 | -0.0214 | 0.0283 | 0.9102 |
| Day 4 | Day 7 | -0.0513 | -0.0015 | 0.0482 | 1.0000 |
| Day 5 | Day 6 | -0.0370 | 0.0127 | 0.0625 | 0.9964 |
| Day 5 | Day 7 | -0.0171 | 0.0326 | 0.0823 | 0.4978 |
| Day 6 | Day 7 | -0.0299 | 0.0199 | 0.0696 | 0.9401 |

**Table S11.** ANOVA results for statistical comparisons between sessions of the maximum depth at K7 and K8.

| Group A | Group B | Lower Limit | A-B | Upper Limit | P-value |
| --- | --- | --- | --- | --- | --- |
| OF1 | OF2 | -0.5654 | 5.3214 | 11.2083 | 0.1104 |
| OF1 | Day 1 | -5.6368 | 0.2500 | 6.1368 | 1.0000 |
| OF1 | Day 2 | -0.5297 | 5.3571 | 11.2440 | 0.1053 |
| OF1 | Day 3 | -2.1725 | 3.7143 | 9.6011 | 0.5505 |
| OF1 | Day 4 | -1.4940 | 4.3929 | 10.2797 | 0.3168 |
| OF1 | Day 5 | -4.9583 | 0.9286 | 6.8154 | 0.9999 |
| OF1 | Day 6 | -4.7797 | 1.1071 | 6.9940 | 0.9996 |
| OF1 | Day 7 | -3.0297 | 2.8571 | 8.7440 | 0.8372 |
| OF2 | Day 1 | -10.9583 | -5.0714 | 0.8154 | 0.1513 |
| OF2 | Day 2 | -5.8511 | 0.0357 | 5.9225 | 1.0000 |
| OF2 | Day 3 | -7.4940 | -1.6071 | 4.2797 | 0.9944 |
| OF2 | Day 4 | -6.8154 | -0.9286 | 4.9583 | 0.9999 |
| OF2 | Day 5 | -10.2797 | -4.3929 | 1.4940 | 0.3168 |
| OF2 | Day 6 | -10.1011 | -4.2143 | 1.6725 | 0.3733 |
| OF2 | Day 7 | -8.3511 | -2.4643 | 3.4225 | 0.9225 |
| Day 1 | Day 2 | -0.7797 | 5.1071 | 10.9940 | 0.1448 |
| Day 1 | Day 3 | -2.4225 | 3.4643 | 9.3511 | 0.6422 |
| Day 1 | Day 4 | -1.7440 | 4.1429 | 10.0297 | 0.3972 |
| Day 1 | Day 5 | -5.2083 | 0.6786 | 6.5654 | 1.0000 |
| Day 1 | Day 6 | -5.0297 | 0.8571 | 6.7440 | 0.9999 |
| Day 1 | Day 7 | -3.2797 | 2.6071 | 8.4940 | 0.8958 |
| Day 2 | Day 3 | -7.5297 | -1.6429 | 4.2440 | 0.9935 |
| Day 2 | Day 4 | -6.8511 | -0.9643 | 4.9225 | 0.9999 |
| Day 2 | Day 5 | -10.3154 | -4.4286 | 1.4583 | 0.3061 |
| Day 2 | Day 6 | -10.1368 | -4.2500 | 1.6368 | 0.3616 |
| Day 2 | Day 7 | -8.3868 | -2.5000 | 3.3868 | 0.9163 |
| Day 3 | Day 4 | -5.2083 | 0.6786 | 6.5654 | 1.0000 |
| Day 3 | Day 5 | -8.6725 | -2.7857 | 3.1011 | 0.8554 |
| Day 3 | Day 6 | -8.4940 | -2.6071 | 3.2797 | 0.8958 |
| Day 3 | Day 7 | -6.7440 | -0.8571 | 5.0297 | 0.9999 |
| Day 4 | Day 5 | -9.3511 | -3.4643 | 2.4225 | 0.6422 |
| Day 4 | Day 6 | -9.1725 | -3.2857 | 2.6011 | 0.7053 |
| Day 4 | Day 7 | -7.4225 | -1.5357 | 4.3511 | 0.9959 |
| Day 5 | Day 6 | -5.7083 | 0.1786 | 6.0654 | 1.0000 |
| Day 5 | Day 7 | -3.9583 | 1.9286 | 7.8154 | 0.9816 |
| Day 6 | Day 7 | -4.1368 | 1.7500 | 7.6368 | 0.9901 |

**Table S12.** Two-sample t-test results for statistical comparisons at each K step of the error rate between real and shuffled data, behaviour-based labels. *\*p*<0.05, *\*\*\*p*<0.001.

| Group A | Group B | h | P-value |
| --- | --- | --- | --- |
| K1 real data | K1 shuffled data | 1 | 3,38E-30*** |
| K2 real data | K2 shuffled data | 1 | 2,82E-30*** |
| K3 real data | K3 shuffled data | 1 | 1,71E-29*** |
| K4 real data | K4 shuffled data | 1 | 4,82E-29*** |
| K5 real data | K5 shuffled data | 1 | 1,39E-28*** |
| K6 real data | K6 shuffled data | 1 | 1,66E-27*** |
| K7 real data | K7 shuffled data | 1 | 3,68E-27*** |
| K8 real data | K8 shuffled data | 1 | 4,12E-25*** |
| K9 real data | K9 shuffled data | 1 | 6,72E-20*** |
| K10 real data | K10 shuffled data | 1 | 2,64E-11*** |
| K11 real data | K11 shuffled data | 1 | 4,01E-02* |

**Table S13.** Statistical comparisons between the error rates at each K step for real data, behaviour-based labels. *\*p*<0.05, *\*\*p*<0.01, *\*\*\*p*<0.001.

| Group A | Group B | Lower Limit | A-B | Upper Limit | P-value |
| --- | --- | --- | --- | --- | --- |
| K1 | K2 | -0,0361 | -0,0005 | 0,0351 | 1 |
| K1 | K3 | -0,0426 | -0,0069 | 0,0287 | 0,9999 |
| K1 | K4 | -0,0498 | -0,0142 | 0,0215 | 0,9725 |
| K1 | K5 | -0,0576 | -0,0220 | 0,0137 | 0,6604 |
| K1 | K6 | -0,0643 | -0,0286 | 0,0070 | 0,2564 |
| K1 | K7 | -0,0715 | -0,0359 | -0,0002 | 0,0467* |
| K1 | K8 | -0,0858 | -0,0501 | -0,0145 | 0,0003*** |
| K1 | K9 | -0,1317 | -0,0960 | -0,0604 | 0,0000*** |
| K1 | K10 | -0,2237 | -0,1881 | -0,1525 | 0,0000*** |
| K1 | K11 | -0,3202 | -0,2584 | -0,1967 | 0,0000*** |
| K2 | K3 | -0,0421 | -0,0064 | 0,0292 | 1 |
| K2 | K4 | -0,0493 | -0,0137 | 0,0220 | 0,9787 |
| K2 | K5 | -0,0571 | -0,0215 | 0,0142 | 0,6915 |
| K2 | K6 | -0,0638 | -0,0281 | 0,0075 | 0,2811 |
| K2 | K7 | -0,0710 | -0,0354 | 0,0003 | 0,0536 |
| K2 | K8 | -0,0853 | -0,0496 | -0,0140 | 0,0004*** |
| K2 | K9 | -0,1312 | -0,0955 | -0,0599 | 0,0000*** |
| K2 | K10 | -0,2232 | -0,1876 | -0,1520 | 0,0000*** |
| K2 | K11 | -0,3197 | -0,2580 | -0,1962 | 0,0000*** |
| K3 | K4 | -0,0429 | -0,0072 | 0,0284 | 0,9999 |
| K3 | K5 | -0,0507 | -0,0150 | 0,0206 | 0,9583 |
| K3 | K6 | -0,0573 | -0,0217 | 0,0140 | 0,6779 |
| K3 | K7 | -0,0646 | -0,0290 | 0,0067 | 0,2409 |
| K3 | K8 | -0,0788 | -0,0432 | -0,0076 | 0,0046** |
| K3 | K9 | -0,1247 | -0,0891 | -0,0534 | 0,0000*** |
| K3 | K10 | -0,2168 | -0,1812 | -0,1455 | 0,0000*** |
| K3 | K11 | -0,3133 | -0,2515 | -0,1898 | 0,0000*** |
| K4 | K5 | -0,0435 | -0,0078 | 0,0278 | 0,9998 |
| K4 | K6 | -0,0501 | -0,0145 | 0,0212 | 0,9680 |
| K4 | K7 | -0,0574 | -0,0217 | 0,0139 | 0,6754 |
| K4 | K8 | -0,0716 | -0,0360 | -0,0003 | 0,0457* |
| K4 | K9 | -0,1175 | -0,0819 | -0,0462 | 0,0000*** |
| K4 | K10 | -0,2096 | -0,1739 | -0,1383 | 0,0000*** |
| K4 | K11 | -0,3060 | -0,2443 | -0,1825 | 0,0000*** |
| K5 | K6 | -0,0423 | -0,0067 | 0,0290 | 1 |
| K5 | K7 | -0,0496 | -0,0139 | 0,0217 | 0,9756 |
| K5 | K8 | -0,0638 | -0,0282 | 0,0075 | 0,2794 |
| K5 | K9 | -0,1097 | -0,0741 | -0,0384 | 0,0000*** |
| K5 | K10 | -0,2018 | -0,1661 | -0,1305 | 0,0000*** |
| K5 | K11 | -0,2982 | -0,2365 | -0,1747 | 0,0000*** |
| K6 | K7 | -0,0429 | -0,0073 | 0,0284 | 0,9999 |
| K6 | K8 | -0,0571 | -0,0215 | 0,0141 | 0,6895 |
| K6 | K9 | -0,1030 | -0,0674 | -0,0317 | 0,0000*** |
| K6 | K10 | -0,1951 | -0,1595 | -0,1238 | 0,0000*** |
| K6 | K11 | -0,2916 | -0,2298 | -0,1681 | 0,0000*** |
| K7 | K8 | -0,0499 | -0,0142 | 0,0214 | 0,9714 |
| K7 | K9 | -0,0958 | -0,0601 | -0,0245 | 0,0000*** |
| K7 | K10 | -0,1879 | -0,1522 | -0,1166 | 0,0000*** |
| K7 | K11 | -0,2843 | -0,2226 | -0,1608 | 0,0000*** |
| K8 | K9 | -0,0815 | -0,0459 | -0,0102 | 0,0017** |
| K8 | K10 | -0,1736 | -0,1380 | -0,1023 | 0,0000*** |
| K8 | K11 | -0,2701 | -0,2083 | -0,1466 | 0,0000*** |
| K9 | K10 | -0,1277 | -0,0921 | -0,0564 | 0,0000*** |
| K9 | K11 | -0,2242 | -0,1624 | -0,1007 | 0,0000*** |
| K10 | K11 | -0,1321 | -0,0703 | -0,0086 | 0,0111* |

**Table S14.** Two-sample t-test results for statistical comparisons at each K step of the maximum tree depth between real and shuffled data, behaviour-based labels. *\*\*\*p*<0.001.

| Group A | Group B | h | P-value |
| --- | --- | --- | --- |
| K1 real data | K1 shuffled data | 1 | 3,79E-04*** |
| K2 real data | K2 shuffled data | 0 | 5,55E-02 |
| K3 real data | K3 shuffled data | 0 | 5,94E-01 |
| K4 real data | K4 shuffled data | 0 | 8,10E-01 |
| K5 real data | K5 shuffled data | 1 | 3,56E-06*** |
| K6 real data | K6 shuffled data | 1 | 1,18E-16*** |
| K7 real data | K7 shuffled data | 1 | 2,56E-26*** |
| K8 real data | K8 shuffled data | 1 | 1,17E-39*** |
| K9 real data | K9 shuffled data | 1 | 1,39E-39*** |
| K10 real data | K10 shuffled data | 1 | 3,56E-27*** |
| K11 real data | K11 shuffled data | 1 | 2,91E-06*** |

**Table S15.** Statistical comparisons between the maximum tree depths at each K step for real data, behaviour-based labels. *\*p*<0.05, *\*\*p*<0.01, *\*\*\*p*<0.001.

| Group A | Group B | Lower Limit | A-B | Upper Limit | P-value |
| --- | --- | --- | --- | --- | --- |
| K1 | K2 | 0,8864 | 7,2000 | 13,5136 | 0,0110* |
| K1 | K3 | 8,7578 | 15,0714 | 21,3850 | 0,0000*** |
| K1 | K4 | 18,1150 | 24,4286 | 30,7422 | 0,0000*** |
| K1 | K5 | 25,5864 | 31,9000 | 38,2136 | 0,0000*** |
| K1 | K6 | 29,5864 | 35,9000 | 42,2136 | 0,0000*** |
| K1 | K7 | 34,5293 | 40,8429 | 47,1565 | 0,0000*** |
| K1 | K8 | 37,0007 | 43,3143 | 49,6279 | 0,0000*** |
| K1 | K9 | 35,7721 | 42,0857 | 48,3993 | 0,0000*** |
| K1 | K10 | 27,9293 | 34,2429 | 40,5565 | 0,0000*** |
| K1 | K11 | 19,6360 | 30,5714 | 41,5069 | 0,0000*** |
| K2 | K3 | 1,5578 | 7,8714 | 14,1850 | 0,0029** |
| K2 | K4 | 10,9150 | 17,2286 | 23,5422 | 0,0000*** |
| K2 | K5 | 18,3864 | 24,7000 | 31,0136 | 0,0000*** |
| K2 | K6 | 22,3864 | 28,7000 | 35,0136 | 0,0000*** |
| K2 | K7 | 27,3293 | 33,6429 | 39,9565 | 0,0000*** |
| K2 | K8 | 29,8007 | 36,1143 | 42,4279 | 0,0000*** |
| K2 | K9 | 28,5721 | 34,8857 | 41,1993 | 0,0000*** |
| K2 | K10 | 20,7293 | 27,0429 | 33,3565 | 0,0000*** |
| K2 | K11 | 12,4360 | 23,3714 | 34,3069 | 0,0000*** |
| K3 | K4 | 3,0435 | 9,3571 | 15,6707 | 0,0001*** |
| K3 | K5 | 10,5150 | 16,8286 | 23,1422 | 0,0000*** |
| K3 | K6 | 14,5150 | 20,8286 | 27,1422 | 0,0000*** |
| K3 | K7 | 19,4578 | 25,7714 | 32,0850 | 0,0000*** |
| K3 | K8 | 21,9293 | 28,2429 | 34,5565 | 0,0000*** |
| K3 | K9 | 20,7007 | 27,0143 | 33,3279 | 0,0000*** |
| K3 | K10 | 12,8578 | 19,1714 | 25,4850 | 0,0000*** |
| K3 | K11 | 4,5645 | 15,7000 | 26,4355 | 0,0003*** |
| K4 | K5 | 1,1578 | 7,4714 | 13,7850 | 0,0065** |
| K4 | K6 | 5,1578 | 11,4714 | 17,7850 | 0,0000*** |
| K4 | K7 | 10,1007 | 16,4143 | 22,7279 | 0,0000*** |
| K4 | K8 | 12,5721 | 18,8857 | 25,1993 | 0,0000*** |
| K4 | K9 | 11,3435 | 17,6571 | 23,9707 | 0,0000*** |
| K4 | K10 | 3,5007 | 9,8143 | 16,1279 | 0,0000*** |
| K4 | K11 | -4,7926 | 6,1429 | 17,0783 | 0,7757 |
| K5 | K6 | -2,3136 | 4,0000 | 10,3136 | 0,6212 |
| K5 | K7 | 2,6293 | 8,9429 | 15,2565 | 0,0003*** |
| K5 | K8 | 5,1007 | 11,4143 | 17,7279 | 0,0000*** |
| K5 | K9 | 3,8721 | 10,1857 | 16,4993 | 0,0000*** |
| K5 | K10 | -3,9707 | 2,3429 | 8,6565 | 0,9833 |
| K5 | K11 | -12,2640 | -1,3286 | 9,6069 | 1 |
| K6 | K7 | -1,3707 | 4,9429 | 11,2565 | 0,2927 |
| K6 | K8 | 1,1007 | 7,4143 | 13,7279 | 0,0073** |
| K6 | K9 | -0,1279 | 6,1857 | 12,4993 | 0,0608 |
| K6 | K10 | -7,9707 | -1,6571 | 4,6565 | 0,9990 |
| K6 | K11 | -16,2640 | -5,3286 | 5,6069 | 0,8957 |
| K7 | K8 | -3,8422 | 2,4714 | 8,7850 | 0,9753 |
| K7 | K9 | -5,0707 | 1,2429 | 7,5565 | 0,9999 |
| K7 | K10 | -12,9136 | -6,6000 | -0,2864 | 0,0316* |
| K7 | K11 | -21,2069 | -10,2714 | 0,6640 | 0,0883 |
| K8 | K9 | -7,5422 | -1,2286 | 5,0850 | 0,9999 |
| K8 | K10 | -15,3850 | -9,0714 | -2,7578 | 0,0002*** |
| K8 | K11 | -23,6783 | -12,7429 | -1,8074 | 0,0082** |
| K9 | K10 | -14,1565 | -7,8429 | -1,5293 | 0,0031** |
| K9 | K11 | -22,4498 | -11,5143 | -0,5788 | 0,0291* |
| K10 | K11 | -14,6069 | -3,6714 | 7,2640 | 0,9922 |

## Notes

### Summary of Updates

Forgot to update the abstract in the prior version, now it is identical to the published version.

